# Biophysical Characterisation of Human LincRNA-p21 Sense and Antisense Alu Inverted Repeats

**DOI:** 10.1101/2021.12.08.471813

**Authors:** Michael H. D’Souza, Tyler Mrozowich, Maulik D. Badmalia, Mitchell Geeraert, Angela Frederickson, Amy Henrickson, Borries Demeler, Michael T. Wolfinger, Trushar R. Patel

## Abstract

Human Long Intergenic Noncoding RNA-p21 (LincRNA-p21) is a regulatory noncoding RNA that plays an important role in promoting apoptosis. LincRNA-p21 is also critical in down-regulating many p53 target genes through its interaction with a p53 repressive complex. The interaction between LincRNA-p21 and the repressive complex is likely dependent on the RNA tertiary structure. Previous studies have determined the two-dimensional secondary structures of the sense and antisense human LincRNA-p21 *AluSx1* IRs using SHAPE. However, there were no insights into its three-dimensional structure. Therefore, we *in vitro* transcribed the sense and antisense regions of LincRNA-p21 *AluSx1* Inverted Repeats (IRs) and performed analytical ultracentrifugation, size exclusion chromatography, light scattering, and small angle X-ray scattering (SAXS) studies. Based on these studies, we determined low-resolution, three-dimensional structures of sense and antisense LincRNA-p21. By adapting previously known two-dimensional information, we calculated their sense and antisense high-resolution models and determined that they agree with the low-resolution structures determined using SAXS. Thus, our integrated approach provides insights into the structure of LincRNA-p21 Alu IRs. Our study also offers a viable pipeline for combining the secondary structure information with biophysical and computational studies to obtain high-resolution atomistic models for long noncoding RNAs.

## 1.0 Introduction

The tumour suppressor protein p53 is an important transcription factor that regulates a variety of cellular processes, including cell-cycle control, DNA repair, apoptosis, senescence, and cellular stress responses through the activation and repression of target genes [1, 2]. Despite playing a critical role in the DNA damage response, p53’s genome is frequently mutated in cancer cells, exposing a vulnerability in cell cycle regulation [3, 4]. Nevertheless, when DNA damage occurs, p53 upregulates the expression of genes involved in the cell cycle arrest and DNA repair processes, which leads to cell survival, but also facilitates the initiation of apoptosis for cancerous cells [5]. The p53 pathway itself is composed of a network of genes, regulatory proteins, and their transcriptional products which can help respond to intrinsic and extrinsic stress signals [8].. These networks enable the regulation of p53 and can be additionally modulated by long noncoding RNAs (LncRNAs) which have been shown to act in a regulatory role within the p53 pathway. Often, the transcription of LncRNA genes are the targets of p53 itself [13–16].

LncRNAs are noncoding RNA molecules devoid of an open reading frame and are generally around 200-100,000 nucleotides (nts). They also do not retain any significant protein-coding capabilities and are therefore generally not expressed [17–19]. LncRNAs were previously thought to have no biological function but have been identified to regulate biological processes by altering gene expression and signal pathways [17]. Consequently, LncRNAs play a role in the regulation of gene expression and appear poised to affect the progression of cancers. Long intergenic noncoding RNA-p21 (LincRNA-p21) is found to be a transcriptional repressor in the p53 pathway, playing a role in triggering cellular apoptosis [20]. LincRNAs are also capped, spliced, and polyadenylated due to being RNA polymerase II transcripts [18]. Under stress conditions including DNA damage, p53 activates the transcription of LincRNA-p21 which accumulates in the nucleus and associates with the heterogeneous nuclear ribonuclear protein K (hnRNP-K) [21]. The hnRNP-K contains RNA recognition motifs Arg-Gly-Gly repeats or hnRNP-K homology (KH) domains and whose role is important for nucleic acid metabolism and transcription [22, 23]. The hnRNP-K is integral in the induction of apoptosis since it will combine with the p53 promoted and transcribed LincRNA-p21 which will then act to repress p53 target genes resulting in apoptosis [24]. LincRNA-p21 is thus required to help direct hnRNP-K to bind to the promoters of the target repressed genes [23]. Additionally, hnRNP-K was observed to be a transcriptional coactivator of p53, enabling gene expression in response to DNA damage [22].

Human LincRNA-p21 exhibits two isoforms which further contains the presence of Alu repeats that which influence the function of the RNA [21]. Alu elements are particularly important because they are highly conserved among primates and fold to produce independent domains. These repeated DNA sequences comprise upwards of 60% of the human genome and can be divided into several classes including micro-satellites (repeat sequences greater than 7 bp), mini satellites (basic repeats of 7 bp or less), or telomeres. These interspersed repeated DNA sequences are further divided into two classes: Short interspersed elements (SINES), and long interspersed elements (LINES) [25]. Alu SINES themselves are repetitive elements present in multiple copies of the genomes they reside in and are named because the family of repeats contains a recognition site for the restriction enzyme *AluI* [26, 27]. Full-length Alu elements are roughly 300 bp long and are frequently located in the 3′-untranslated regions of genes and their intergenic genomic regions and continue to be the most abundant mobile or transposable element in the entirety of the human genome. Determining the structural-dependent role of LincRNA-p21 Alu elements will have an impact on elucidating their overall function and responsibilities within the cell.

Many studies using molecular and computational structural biology seek to identify LncRNA secondary and tertiary structures, and whether said structures have an impact on their function [31, 32]. This also includes the application of RNA secondary structure prediction techniques such as selective 2′-hydroxyl acylation analysed by primer extension (SHAPE) [33]. Doing so is important because it conceptualises structures present on LincRNA-p21 and can elucidate potential specific interactions within the p53 and hnRNP-K pathways. Previous studies have investigated the secondary structure of LincRNA-p21 Alu Inverted Repeats (IRs) and identifying important functional regions that are involved in LincRNA-p21 nuclear localisation and its subsequent transcriptional factor interactions [21]. They identified that the two isoforms of LincRNA-p21 contained Alu IRs that retained integral secondary structures that can fold into independent domains. These structures were suggested to be conserved in primates and contribute towards the regulation of cellular localisation of LincRNA-p21 during the cellular stress response.

Incidentally, the intent of this study is to investigate the overall structure of the sense and antisense LincRNA-p21 *AluSx1* Inverted Repeats by employing small angle X-ray scattering (SAXS) and computational modelling to develop their three-dimensional structures [34]. We have employed analytical ultracentrifugation (AUC) and size exclusion chromatography coupled to multiangle light scattering (SEC-MALS) instruments to biophysically characterise transcribed and purified *AluSx1* RNAs. AUC experiments revealed that *AluSx1* RNAs were present as monomeric, full-length transcripts under denaturing conditions, while SEC-MALS characterised their Molecular Weight (MW). By combining chemically probed secondary structure information proposed by Chillón and Pyle, 2016, and SimRNA computational modelling, several three-dimensional, high-resolution models can be calculated and fitted to SAXS determined structures. We determined that human LincRNA-p21 isoform LIsoE2’s *AluSx1* RNA adopts an asymmetrical, and extended structure in solution. The importance being that no previous three-dimensional structure determination has been performed on LincRNA-p21 *AluSx1* IRs which can improve on the understanding of its interactions with hnRNP-K. We describe a workflow utilising SAXS and SimRNA computational modelling to produce three-dimensional, high-resolution models devised from two-dimensional structures determined *via* SHAPE and other secondary structure probing techniques.

## 2.0 Methods

### 2.1 Sense and Antisense LincRNA-p21 *AluSx1* Plasmid Preparation

A flowchart of the procedure is outlined in **Figure 1**. LincRNA-p21 *AluSx1* transcripts for the sense and antisense (taken from the TP53COR1 gene located on Chr6:36,663,392-36,667,296 (GRCh38/hg38), Chr6:36,631,169-36,635,073 (GRCh37/hg19)) were designed from the hLincRNA-p21 LIsoE2 isoform sequences presented in from Chillón and Pyle [20, 21]. *AluSx1* IRs are bidirectionally transcribed from the total TP53COR1 gene, producing sense and antisense designated transcripts [21]. Thus, RNA constructs used in this experiment are represented below:

>LincRNA-p21 *AluSx1* Sense RNA Sequence (307nt) | TP53COR1_LIsoE2_*AluSx1*_P

5′-

AGCUGGGCGUGGUGGCUCACGCCUGUAAUCCCACCACUUUGGGAGGCCGAGGCAGGCGGAUCACUUGAGGUCAGGAGUCCAAGACCAGCCUGGCCAACAAGGCGAAACCCUGUCUCUACUAAAAAUACAAAAACUAGCUGGGCGUAGUGGUGGGCACCUGUAAUCCCAGCUACUCGGGAGGCUGAGACAGGACAAUCGCUUGGACUCCGGAGGCAGAGGUUGCAGUGAGCUGGGAUCGUGCCACUACACUCCAGUCUGGGCGACAGAGCAAGACUCUGCAUCAAAAAAAAAAAAGAAAGAGUAAUAA-3′

>LincRNA-p21 *AluSx1* Antisense RNA Sequence (280nt) | TP53COR1_LIsoE2_*AluSx1*_P

5′-

GCAGAGGAGGAAUGGAAUCAUUCUUUUUUUUUUUAUUGGAGACGGAGUCUCACUCUGUUGCUCAGGCUGGAGUGUAGUGGUGCGAACUUGGCUCACUGCAGCCUCCACCUCCCAGGCUCAAGCAAUUCUCCUGCCUCAGCCUCCCGAGUAGCUGGGAUUACAGGUGUCUGCUAUCACACCCAGCUAAAGUUUUUAUAUUUUUAGUAGAAAUGGAGUUUCACCAUGUUGGACAGGCUGGUCUCGAACUCCUGACCUCAGGUGAUCCACCCGCCUCAGCCUC-3′

**Figure 1:**
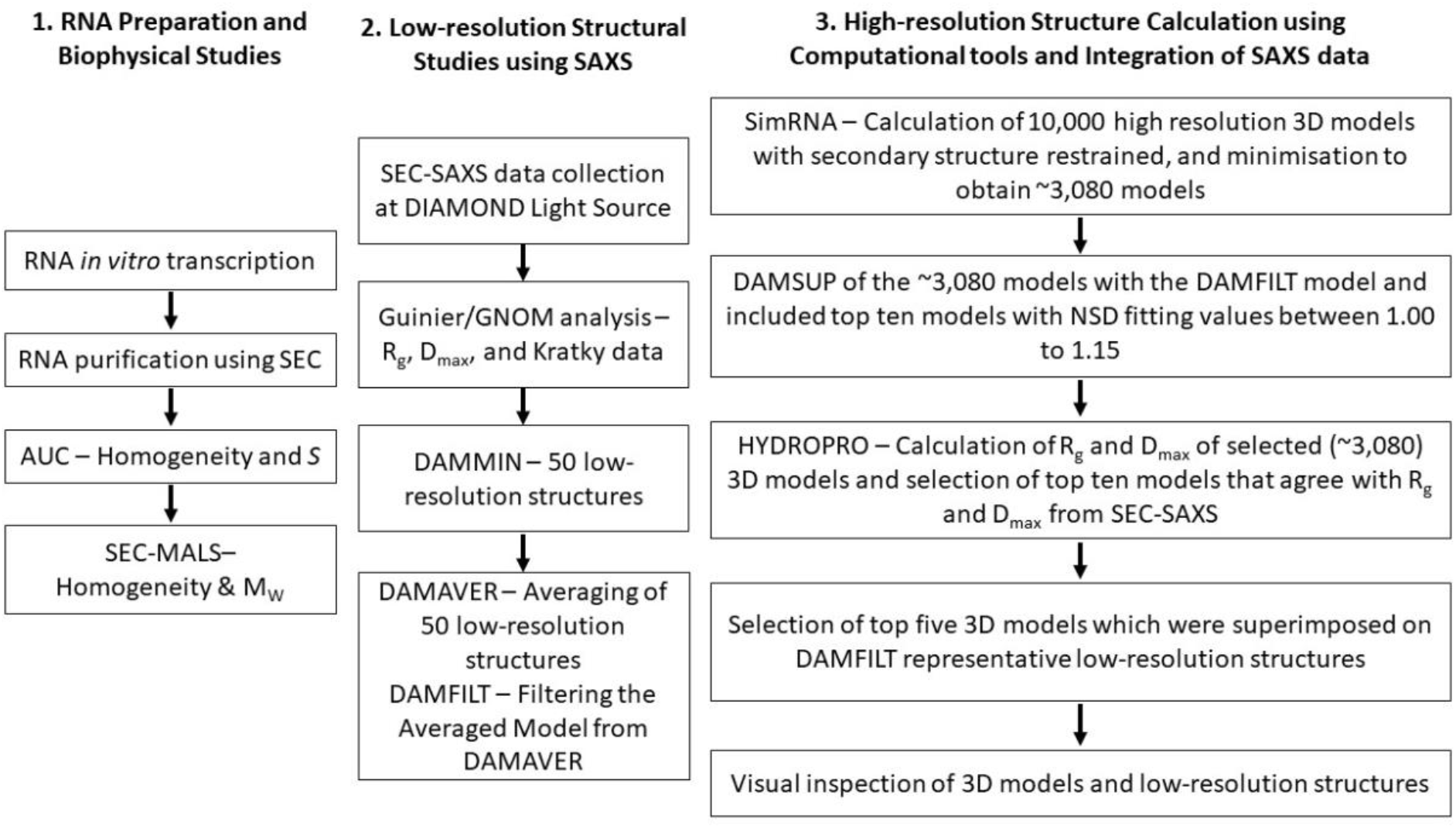
Organisational Flowchart for the Purification and Characterisation of Sense and Antisense LincRNA-p21 *AluSx1* RNA. The determination of LincRNA three-dimensional, low-resolution structures overlaid by high-resolution, atomistic models was conducted in three phases: RNA preparation and biophysical studies to determine sample homogeneity and sample properties; low-resolution structure determination by SAXS; and high-resolution modelling using SimRNA, with constraints imposed by HYDROPRO. All methods are further described below.

This human LincRNA-p21 isoform LIsoE2 *AluSx1* RNAs will be referred to as sense and antisense *AluSx1* RNAs throughout.

The plasmids were synthesised commercially, each sequence was flanked by a T7 RNA polymerase promoter sequence at the 5′-end and an *XbaI* restriction endonuclease cut-site sequence at the 3′-end. To increase RNA yield, two additional Gs were added to the 3′-end of the T7 promoter region which is reflected in the theoretical MW (**Table 1**) [35]. LincRNA-p21 sequences were inserted into Genewiz pUC-57-KAN plasmids (Azenta Life Sciences, USA). Plasmids were transformed and cultured in *E. coli* NEBα (NEB, Canada) competent cells and were purified using NEB Monarch Miniprep Kits (NEB, Canada) as per manufacturer’s protocol.

**Table 1:** Solution Properties of Sense and Antisense LincRNA-p21 *AluSx1*.

| Sample | Sense LincRNA-p21 *AluSx1* | Antisense LincRNA-p21 *AluSx1* |
| --- | --- | --- |
| $M_w$ Theoretical (kDa) | 99.418 | 89.543 |
| $M_w$ AUC (kDa) <sup>+</sup> | 94.770 | 92.561 |
| $M_w$ SEC-MALS (kDa) <sup>∇</sup> | 99.24 ± 0.01 | 94.52 ± 3.71 |
| Sedimentation Coefficient, $s_{20,w}$ ($10^{-13}$ s) <sup>+</sup> | 5.56 ± 0.25 | 5.53 ± 0.05 |
| Diffusion Coefficient, $D_{20,w}$ ($10^{-7}$ cm<sup>2</sup>/s) <sup>+</sup> | 2.95 ± 0.59 | 3.00 ± 0.03 |
| Frictional Ratio, $f/f_0$ <sup>+</sup> | 2.71 | 2.68 |
| $R_h$ (Å) <sup>+</sup> | 72.7 | 71.3 |
| $q.R_g$ range <sup>#</sup> | 0.43-1.29 | 0.42-1.25 |
| $R_g$ (Å) <sup>#</sup> | 60.87 ± 0.85 | 59.07 ± 0.15 |
| $I(0)$ <sup>Δ</sup> | 0.01 ± 9.90 × 10<sup>-5</sup> | 0.07 ± 8.14 × 10<sup>-5</sup> |
| $R_g$ (Å) <sup>Δ</sup> | 61.71 ± 0.31 | 58.37 ± 0.07 |
| $D_{max}$ (Å) <sup>Δ</sup> | 185.0 | 180.7 |
| $X^2$ <sup>*</sup> | ~1.148 | ~1.084 |
| NSD <sup>*</sup> | 1.080 ± 0.024 | 1.005 ± 0.022 |

The $M_w$ of the sense and antisense LincRNA-p21 *Alu* Repeat RNA were calculated using the nucleotide sequences provided by Chillón and Pyle, 2016. <sup>+</sup> Molar masses and frictional ratios determined by AUC assume a partial specific volume of 0.516 mL/g and refer to conditions where the RNA is denatured by 6M urea. <sup>+</sup> are within 95% confidence intervals. Data points <sup>∇</sup> were determined from SEC-MALS experiments. Data points <sup>#</sup> were derived from the Guinier analysis. Data points <sup>Δ</sup> were determined using $P(r)$ analysis using the GNOM program. Data points <sup>\*</sup> were derived from DAMMIN and DAMAVER analysis. Terms: Hydrodynamic Radius ( $R_h$ ); Radius of Gyration ( $R_g$ ); Maximum Particle Dimension ( $D_{max}$ ); Normalised Spatial Discrepancy (NSD).

### 2.2 *In vitro* Transcriptions of LincRNA-p21 Sense and Antisense *AluSx1* Inverted Repeats and RNA Purification

RNA transcripts were prepared using run-off *in vitro* transcriptions (IVT) as prepared previously [36, 37]. Briefly, concentrated plasmid samples were digested by *XbaI* restriction endonuclease (NEB, Canada). 1 mL *in vitro* transcription reactions were performed using laboratory purified in-house T7 RNA polymerase and commercial RiboLock RNase inhibitor (Thermo Fisher Scientific, USA) [38]. Linearised plasmids were additionally incubated with 10% DMSO and 0.1% Triton X-100 to increase RNA transcript yields [39]. Sense and antisense *AluSx1* RNA were purified by SEC using a Superdex 200 Increase GL 10/300 (Global Life Science Solutions USA LLC, Marlborough, MA, USA) and purification buffer (PB) (10 mM Bis-tris pH 5.0, 100 mM NaCl, 15 mM KCl 15 mM MgCl2, 10% glycerol) with an ÄKTA pure FPLC (Global Life Science Solutions USA LLC, Marlborough, MA, USA) [40]. SEC peak fractions were assessed for purity by urea-polyacrylamide gel electrophoresis (Urea-PAGE) and sedimentation velocity analytical ultracentrifugation (SV-AUC) in 6M urea. Urea-PAGE (10%) was run at room temperature, 300V, for 40 min in 1x TBE (Tris-Borate-EDTA) buffer, followed by staining with Sybr™ Safe (Thermofisher Scientific, Saint-Laurant, QC, Canada) and visualisation. Pure fractions were pooled and concentrated by ethanol precipitation, with resuspension in HEPES Folding Buffer (HFB) (50mM HEPES, 150 mM NaCl, 15 mM MgCl2, 3% Glycerol, pH 7.4) for SAXS submission.

### 2.3 Multiangle Light Scattering (MALS), and Analytical Ultracentrifugation (AUC) Studies of LincRNA-p21 *AluSx1* Sense and Antisense Inverted Repeats

SEC purified LincRNA-p21 *AluSx1* RNAs were subjected to an additional SEC purification by a Superdex 200 Increase 10/300 GL column in HFB and analysed directly by an in tandem DAWN Multiangle Light Scatterer (MALS) with Optilab Refractive Index System (Wyatt Technology, USA) to determine the MW as per Wyatt Technologies guidelines [41]. Samples were eluted at a 0.5 mL/min flowrate and measured using 18 multiangle detectors, including a UV A_260_ and A_280_, and a refractive index (RI) detector. MALS measurements were taken using a helium-neon red laser (632.8 nm) at 25°C. For data analysis, the refractive index increment (dn/dc) was adjusted to 0.1721 mL/g for sense and antisense *AluSx1* RNA samples [42–44]. Data were analysed using the ASTRA v9 software package (Wyatt Technology, USA) [45, 46].

Purified sense and antisense *AluSx1* RNA were measured by SV-AUC under denaturing conditions in 6M urea to ascertain purity and composition of the transcript. Both samples were measured in two-channel centrepieces and spun at 25,000 rpm for 6 hours at 20°C. Denaturing 6M urea buffer density (1.0899 g/mL) and viscosity (1.3896 cP) were estimated with Ultrascan and used to convert observed sedimentation and diffusion coefficients to standard conditions (water at 20°C). Data were collected in intensity mode at 260 nm and processed using the UltraScan III Software [47]. SV-AUC data were processed as described in [48]. Briefly, systematic noise contributions and boundary conditions (meniscus and bottom of the cell position) samples were processed with the two-dimensional spectrum analysis [49]. Data was further refined by genetic algorithm analysis to achieve parsimonious regularisation [50]. The final step included a genetic algorithm-Monte Carlo (GA-MC) analysis, that was performed with 50 iterations to obtain 95% confidence intervals for the determined parameters (**Table 1**) [51]. AUC data analysis was performed on the XSEDE high-performance computing infrastructure using Expanse and Bridges-2 at the San Diego and Pittsburgh supercomputing centres, respectively. The final model produced very low residual mean square deviations (RMSD) of 0.00139 at 0.438 OD_260_ for sense and 0.00177 at 0.71 OD_260_ for antisense LincRNA-p21 *AluSx1*. All fits produced random residuals, which, together with the low RMSD is evidence for excellent convergence.

### 2.4 Small Angle X-Ray Scattering (SAXS) Analysis of LincRNA-p21 *AluSx1* Sense and Antisense

SAXS data for sense and antisense *AluSx1* RNA samples were collected at 2.5 mg/mL. Samples were run at Diamond Light Source Ltd. synchrotron (Didcot, Oxfordshire, UK) on the B21 SAXS beamline, with a high-performance liquid chromatography (HPLC) system attached upstream to ensure sample monodispersity [52]. A specialised flow cell was employed in conjunction with an inline Agilent 1200 HPLC system (Agilent Technologies, Stockport, UK); sense and antisense *AluSx1* RNA samples were injected onto a Shodex KW403-4F (Showa Denko America Inc., New York, NY, USA) size exclusion column pre-equilibrated with HFB. The flow rate of the column was maintained at 0.160 mL/minute with eluted samples being exposed to X-rays with 3 second exposure time and 600 frames.

Analysis of scattering data was carried out using the ATSAS suite [53]. Using Chromixs, the buffer contribution was subtracted from the sample peak [54]. A Guinier analyses (q^2^ vs. ln(I(q))) was performed on each data set to obtain the radius of gyration (R_g_) and to determine the sample’s quality [55]. A dimensionless Kratky plot (qR_g_ vs qR_g_^2∗^I(q)/I(0)) was generated to evaluate folding of RNA molecules [56]. A paired-distance distribution function (P(r) analysis was performed using GNOM to obtain real-space Rg and the maximum particle dimension (D_max_) of the sample [57, 58]. Employing the information derived from the P(r) plot, a total of fifty sense and antisense *AluSx1* RNA models were generated using DAMMIN [59]. These models were then averaged using DAMAVER and then filtered using DAMFILT to produce a single representative model of each of the RNAs [59, 60].

### 2.5 Sense and Antisense LincRNAp-21 AluSx1 RNA Tertiary Structure Determination

Using the secondary structure information from Chillón and Pyle, 2016, sense and antisense *AluSx1* tertiary structures were calculated using SimRNA v3.20 [34]. SimRNA v3.20 is a Monte Carlo sampler that operates on a coarse-grained model of RNA structure. SimRNA employs a five-bead system per nucleotide, as well as an empirically derived knowledge-based potential. A total of 20 million SimRNA iterations in replica exchange mode were performed for both sense and antisense *AluSx1* RNAs. SimRNA clustering was then performed within one percent of all trajectories with the lowest energy. A RMSD cut-off of five was applied to filter 3080 clusters of similar structures for both sense and antisense *AluSx1* RNAs.

### 2.6 High-Resolution Structural Modelling of LincRNA-p21 *AluSx1* Sense and Antisense

The representative cluster models containing 3080 computationally generated high-resolution models for both the sense and antisense *AluSx1* were separately assessed by HYDROPRO to generate hydrodynamic properties for each model [61]. Running conditions for HYDROPRO involved buffer properties for HFB as determined by UltraScan III: a viscosity of 1.10068 cP; and buffer density of 1.014 g/cm^3^ [47]. The theoretical MW of 99.418 kDa and 89.543 kDa were applied to the HYDROPRO parameters of sense and antisense *AluSx1* RNA, respectively. Models were superimposed onto the SAXS DAMFILT structures and fitted using DAMSUP. Models exhibiting an NSD (normalized spatial discrepancy) value of 1.00 to 1.15 which indicates close fitting were further selected to represent the high-resolution, atomistic RNA model [60]. Models exhibiting similar HYDROPRO determined R_g_ and D_max_ were further selected for and formed the top ten models of interest. The top ten models were energy minimised using an additional step involving QRNAS, which employed a subset of the AMBER force field to achieve energy minimisation of the structures generated from coarse-grained three-dimensional modelling [62]. 20,000 QRNAS MD iterations were performed from the original SimRNA full-atom reconstructed high-resolution models that best-fit the averaged, filtered low-resolution, three-dimensional structure obtained from DAMFILT. Subsequently, five best fit models were superimposed on SAXS structures and represented using PyMOL [60, 63].

## 3.0 Results

### 3.1 Purification of LincRNA-p21 Sense and Antisense *AluSx1* Inverted Repeats

Both sense and antisense *AluSx1* RNAs were purified using SEC, eluting principally around ~10.0 – 12.5 mL at a flowrate of 0.5 mL/min on the Superdex 200 Increase GL 10/300 (**Figure 2A**). The left peak indicates plasmid excluded from subsequent analysis while the right peak represents the RNA of interest. **Figure 2B** depicts the 10% Urea PAGE gel for the RNA fractions indicating that both RNAs migrated closely with similar length and, around the ~300bp marker. Sense LincRNA-p21 *AluSx1* generally produced closely eluting bands (around ~300bp). SV-AUC experiments were conducted using the single fractions collected at 11.0 mL and 11.5 mL for sense and antisense *AluSx1* RNA respectively (indicated by the right shoulder, blue inset, **Figure 2A**).

**Figure 2:**
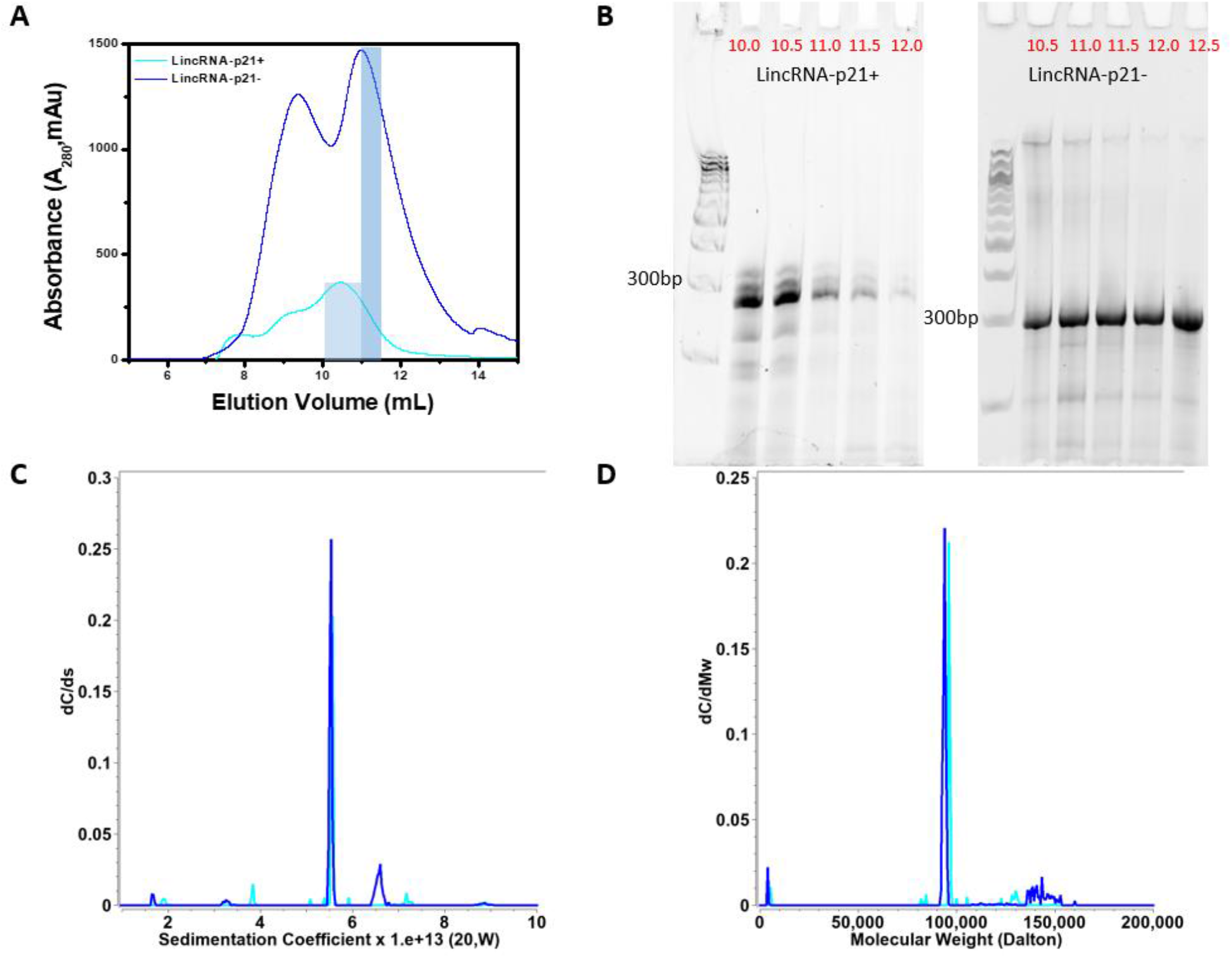
Purification of Sense and Antisense *in vitro* Transcribed LincRNA-p21 *AluSx1* RNA. (**A**) depicts the size exclusion chromatogram of the sense and antisense *AluSx1* RNA elution profile using the Superdex 200 Increase GL 10/300 column. SEC-MALS and SV-AUC experiments were performed with the fractions highlighted in light blue (sense) and dark blue (antisense). (**B**) shows the 10% urea PAGE gel used to ascertain the sense and antisense LincRNA-p21 RNA purity extracted using 0.5 mL fractions (volumes in red) using an ÄKTA Pure FPLC through a Superdex 200 Increase GL 10/300 SEC column. Fractions collected at 11.0 mL and 11.5 mL for sense and antisense *AluSx1* purifications were consolidated and used for SAXS and SV-AUC experiments. A Quick-Load® Purple 100 bp DNA Ladder (NEB, Canada) was used for the 10% urea PAGE gels in lanes 1 and 7 of each gel. (**C**) dC/ds sedimentation coefficient distributions for sense (light blue) and anti-sense (dark blue) under 6M urea denaturing conditions. (**D**) same as (**C**), except transformed to molar mass distributions assuming a partial specific volume of 0.516 mL/g.

### 3.2 Biophysical Characterisation of LincRNA-p21 Sense and Antisense AluSx1 Inverted Repeats

Sedimentation and diffusion coefficients and their 95% confidence intervals resulting from the GA-MC analyses are listed in **Table 1**. Together with sequence based molar masses, SV-AUC results can be used to derive partial specific volumes and anisotropies for the RNA measurements. Since both RNA molecules were measured in the same urea buffer, it is reasonable to assume that the partial specific volume is similar for both molecules. Sedimentation experiments were performed in 6M urea to denature the molecule and disrupt hydrogen bonding within double-stranded RNA regions of the molecule. Results shown in **Figure 2C** and **Figure 2D** indicate that both samples contain one major species with similar sedimentation coefficients, and molar masses in agreement with molar masses predicted from sequence when using a partial specific volume of 0.516 mL/g. This result is consistent with a monomeric and homogeneous full-length transcript of sense (84% of total concentration) and antisense (75% of total concentration) LincRNA-p21 *AluSx1* RNA. Frictional ratios and hydrodynamic radii derived for both molecules indicate a high anisotropy for both molecules, consistent with an unfolded and extended molecule.

SEC-MALS analysis was conducted to determine the MW of the sense and antisense *AluSx1* RNA SAXS. LincRNA-p21 *AluSx1* RNAs were purified again on a Superdex 200 Increase GL 10/300 SEC column which produced peaks eluting between 11 – 12.5 mL (**Figure 3A**). MW values of sense and antisense *AluSx1* reported from SEC-MALS are slightly higher than the molar masses calculated from their sequences except for sense LincRNA-p21 *AluSx1* which exhibited less than 0.2% difference from the theoretical MW at 99.24 ± 0.01 kDa (**Table 1**). MW uniformity throughout **Figure 3B** and **Figure 3C** indicates that both RNAs are monomeric. SEC-MALS results further confirm the homogeneous composition of sense and antisense *AluSx1* RNA determined in SV-AUC. RNA degradation or shorter transcripts can be excluded since no significant smaller fragments were detected. Hydrodynamic parameters derived from SEC-MALS, and SV-AUC are summarised in **Table 1**.

**Figure 3:**
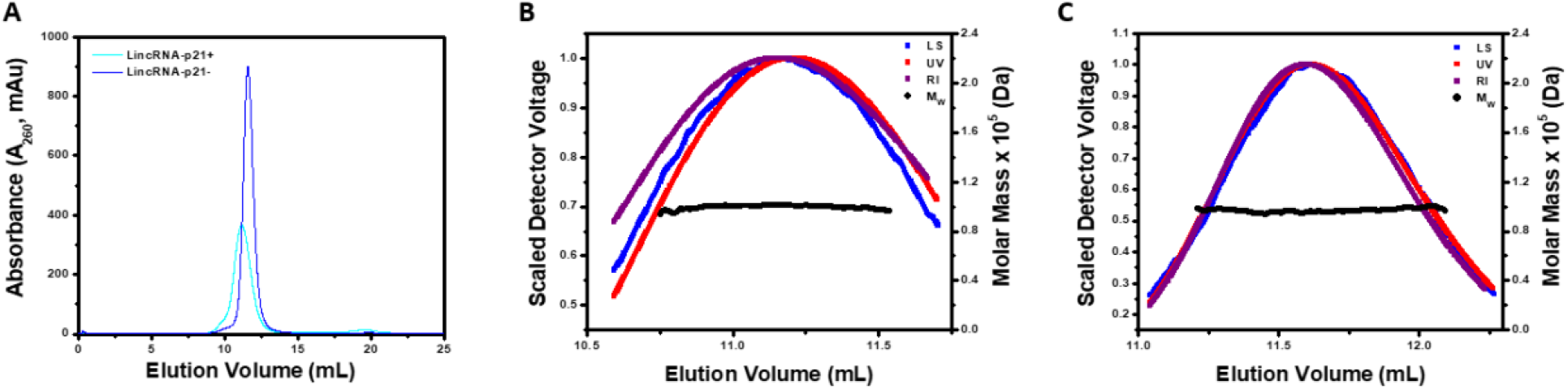
Molecular Weight Determination of Sense and Antisense LincRNA-p21 *AluSx1* RNA using SEC-MALS. (**A**) Portrays the elution curve from the Superdex 200 Increase GL 10/300 SEC of sense and antisense *AluSx1* RNAs. (**B**) Demonstrates the absolute molecular weight distribution across the elution peak of sense LincRNA-p21 *AluSx1* RNA’s elution profile, and light scattering (blue), UV (red), and RI (purple) scattering. (**C**) Portrays the absolute molecular weight distribution across the elution peak the results fitting of antisense LincRNA-p21 *AluSx1* RNA’s elution profile, and light scattering (blue), UV (red), and RI (purple) scattering.

### 3.3 Low-Resolution Structural Studies of LincRNA-p21 Sense and Antisense *AluSx1* Inverted Repeats

SAXS is a powerful method that can represent the overall solution shape of biomolecules under physiologically relevant conditions. Using SEC-SAXS, which can separate different species according to their size before being applied to the SAXS measuring cell, provides confidence in the monodispersity of purified samples [64–67]. The resulting datasets were merged and presented in **Figure 4A** depicting the scattering intensity relative to angle for sense and antisense *AluSx1* RNA. A Guinier analysis (l(q)) vs. (q^2^)) represented by **Figure 4B** displays the LincRNA-p21 *AluSx1* RNA samples’ purity [55]. The Guinier analysis determined the Guinier Rg from the low-q region as being 60.87 ± 0.87 Å for sense LincRNA-p21 *AluSx1* and 59.07 ± 0.15 Å for antisense LincRNA-p21 *AluSx1*. Intensity data from **Figure 4A** was transformed to produce a dimensionless Kratky plot (**Figure 4C**) to determine the LincRNA-p21 *AluSx1* RNAs’ conformations in solution [68]. The dimensionless Kratky plot for both the sense and antisense AluSx1 shows a levelled-plateau which suggests them as being folded and extended in solution [69].

**Figure 4:**
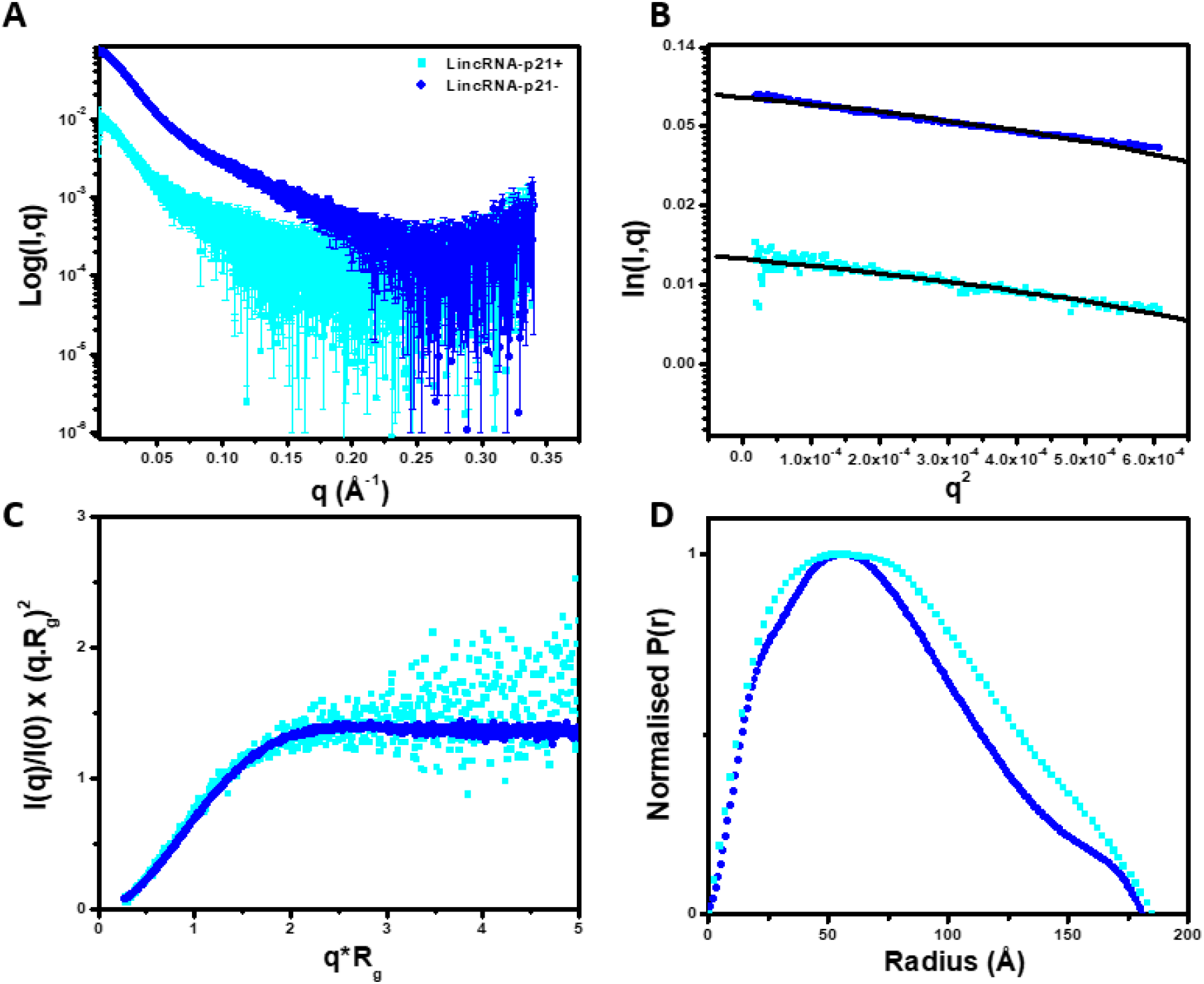
Small Angle X-Ray Scattering (SAXS) Characterisation of Sense and Antisense LincRNA-p21 *AluSx1* RNA. (**A**) merged scattering data of sense and antisense *AluSx1* RNA depicting the scattering intensity (log I(q)) vs. scattering angle (q = 4πsinθ/λ). (**B**) Guinier plots allowing for the determination of R_g_ from the low-angle region data and representing the homogeneity of samples. (**C**) Dimensionless Kratky plots (I(q)/I(0)*(q*R_g_)2 vs. q*R_g_) of sense and antisense *AluSx1* RNA depicting the elongated, tube-like structures because of the non-Gaussian, levelled-plateau shape of the curve. (**D**) Normalised pair distance distribution plots for sense and antisense *AluSx1* RNA which permits the determination of R_g_ derived from the SAXS dataset and including each molecule’s D_max_.

**Figure 4D** represents the (P(r)) plot which was derived from indirect Fourier transformations to convert the reciprocal-space information of the intensity data in **Figure 4A** to real-space electron pair distance distribution data [70]. Using the P(r) plot, sense LincRNA-p21 *AluSx1* presented a real-space R_g_ of 61.71 ± 0.31 Å and a D_max_ of 185.0 Å, while the antisense LincRNA-p21 *AluSx1* presented a real-space R_g_ of 58.37 ± 0.07 Å and D_max_ of 180.7 Å. DAMMIN was performed to obtain low-resolution structures for the sense and antisense *AluSx1* RNA. Fifty models were calculated for each sense and antisense *AluSx1* RNAs which demonstrated favourable agreement as indicated by the X^2^ values (~1.148 for sense LincRNA-p21 *AluSx1* and ~1.084 for antisense LincRNA-p21 *AluSx1*). DAMFILT and DAMAVER were performed to filter and averaged the models. The NSDs were estimated to be 1.080 ± 0.024 and 1.005 ± 0.022 for sense and antisense respectively (**Table 1**) [63].

We identified two, single representative SAXS envelopes illustrated by **Figure 5**. The averaged, single-representative SAXS envelope of sense LincRNA-p21 *AluSx1* is generally extended, adopting a non-spherical, nonglobular surface model (**Figure 5A**). The SAXS envelope is additionally asymmetrical in its rotation along its x- and y-axes, exhibiting two prominent bulges that are primarily located on its ends. **Figure 5B** shows the antisense LincRNA-p21 *AluSx1* SAXS envelope which is similarly elongated and asymmetrical. Antisense LincRNA-p21 *AluSx1* though has three prominent bulges, two located centrally, while the third distally protrudes outwards from the centre.

**Figure 5:**
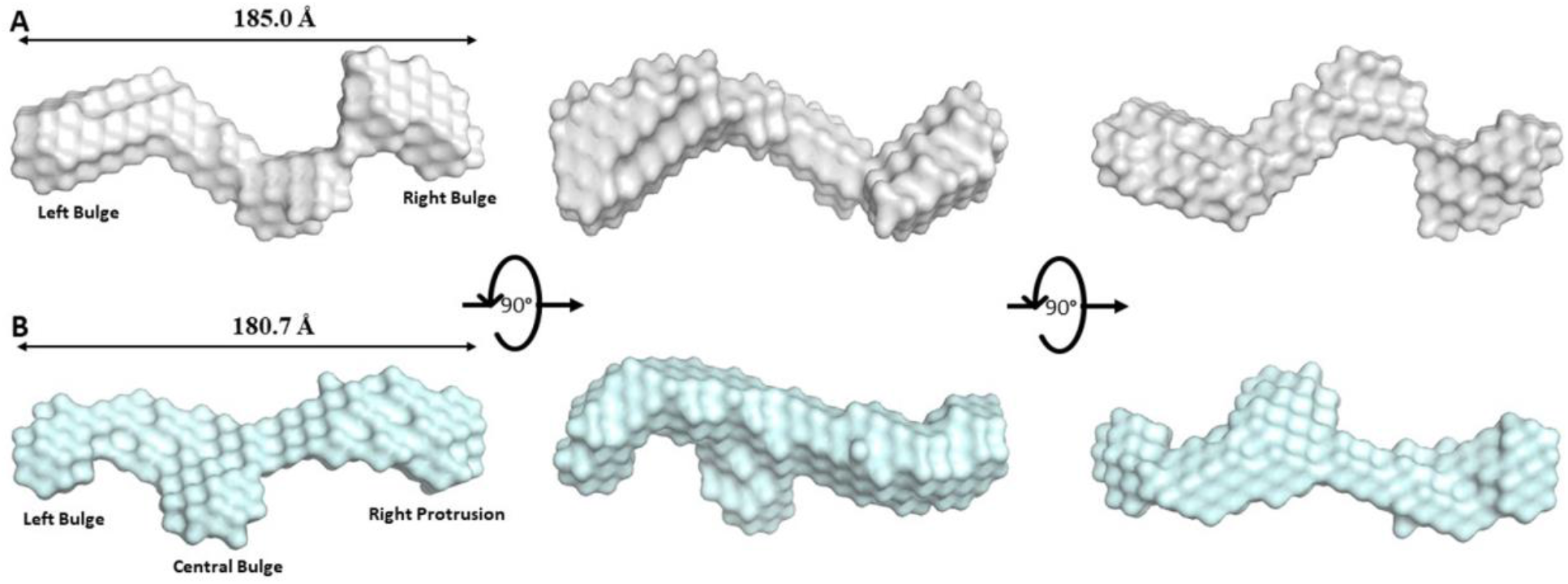
Low-Resolution Structures of Sense (A, Grey) and Antisense (B, Pale Cyan) LincRNA-p21 *AluSx1* Inverted Repeats Determined using SAXS. **(A)** The averaged DAMAVER SAXS low-resolution structure of sense LincRNA-p21 *AluSx1* RNA, taking on an elongated, asymmetrical, and extended structure with maximum length of 185.0 Å. Key features include a left and right Bulge. **(B)** The averaged DAMAVER SAXS low-resolution structure of antisense LincRNA-p21 *AluSx1* RNA, adopting an elongated, asymmetrical, and extended structure with maximum length of 180.7 Å. Key features include a left bulge, central bulge, and a right protrusion. Dimensions are represented by the D_max_ obtained from the P(r) analysis. Models are rotated along their x-axis by 90° as represented by the inset.

### 4.2.4 High-Resolution Atomistic Models of LincRNA-p21 *AluSx1* Sense and Antisense Inverted Repeats

Using SimRNA v3.20, and the secondary structure constraints for both RNAs based on previous studies, we calculated 10,000 clusters of high-resolution, atomistic models for each RNA [21]. These models were further refined through energy minimisation steps to remove models that did not satisfy constraints such as defined atom distances, bond lengths and angles. Subsequently, we obtained 3080 high-resolution models that can be superimposed on the DAMFILT SAXS envelopes using DAMSUP. Sense LincRNA-p21 *AluSx1* models when superimposed produced ten models that had an NSD range from 1.053 to 1.094, while for the antisense RNA, the top ten models retained an NSD range of 1.113 to 1.175. We further applied a selection process using the real-space R_g_ and D_max_ values determined from SAXS for each molecule. We employed the program HYDROPRO to calculate biophysical properties such as R_g_ and D_max_ from the 3080 high-resolution models, as performed previously [71]. Top ten models were further reduced to five using the HYDROPRO properties to achieve models that were in close approximations to SAXS determined R_g_ and D_max_.

Both the top five high-resolution, high-fidelity sense and antisense *AluSx1* models are represented by **Figures 6** and **7** respectively. **Figure 6** depicts the sense LincRNA-p21 *AluSx1* RNAs that closely fit with the SAXS envelopes generated in **Figure 5A**. Previous chemically probed secondary structure predictions identified three major secondary structures: a left and right arm, and a 3′-three-way junction which have been modelled using SimRNA and represented in **Figure 6**. High-fidelity sense LincRNA-p21 *AluSx1* characteristically exhibits the right arm (Magenta) that curls into the central RNA body while the left arm (Blue) extends outwards. Multiple models depict variance in the right arm’s position, appearing to adopt multiple conformations that curl, but rarely extend outwards, towards the central RNA body. The right arm’s stem-loop on its head is additionally compacted and has either bridged with the main RNA body against the 3′-three-way junction or curls outwards. We have identified a consistent 3′-adenyl tail (Cyan) that consistently wraps around the right arm’s base or the connection with the 3′-three-way junctions (Yellow). A 5′-junction (Green) is also a present feature identified but not named by the previous study, but consistently appears to project outwards, perpendicularly from the RNA’s x-axis (**Figure 6**). A flexible, and generally unnamed region – the single-stranded linker (Orange), is presented centrally between the two arms, adopting no specific structure. An animated representation of **Figure 6B** is attached in **Supplementary Information** as **SM 1**.

**Figure 6:**
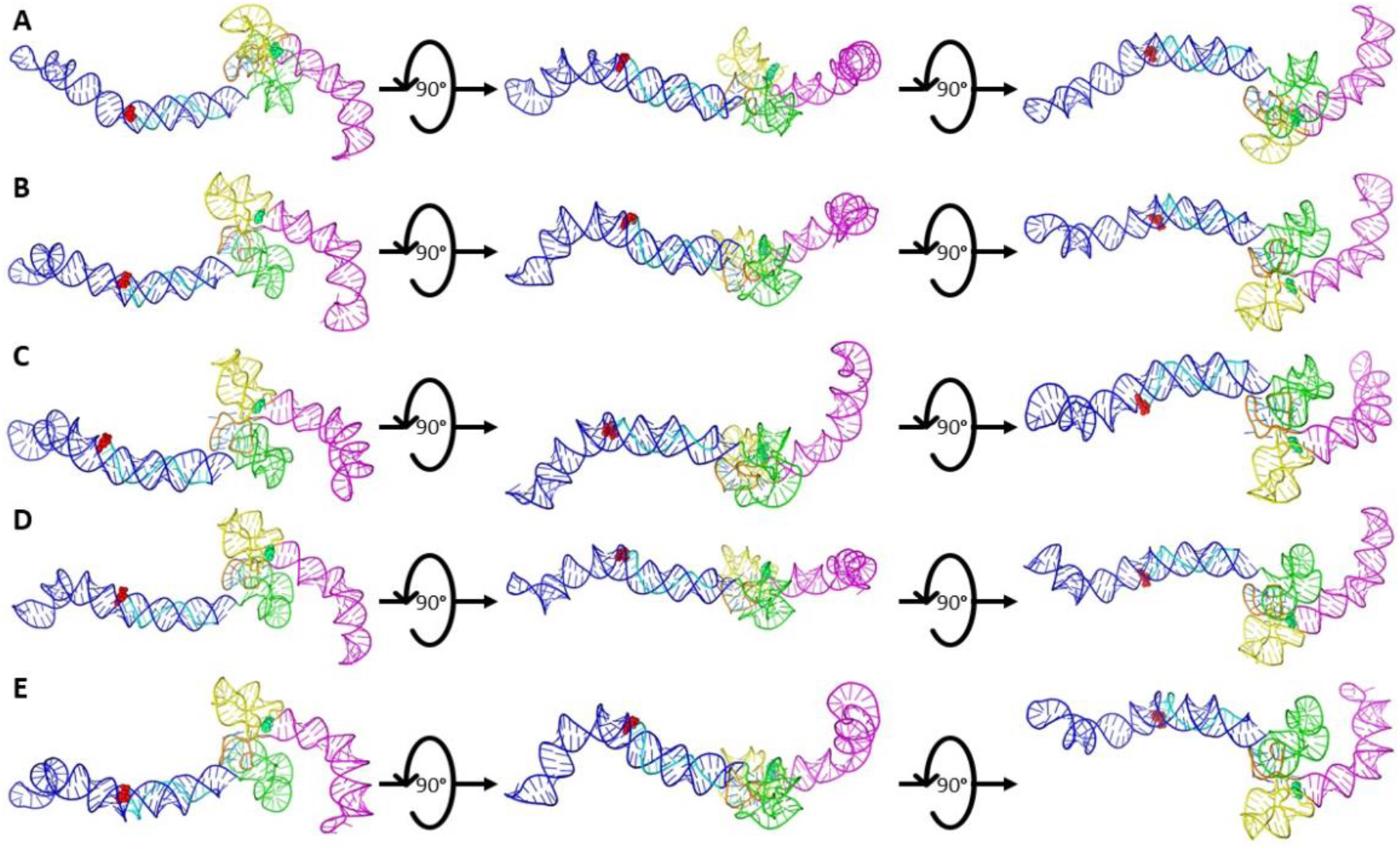
The SimRNA High-Resolution, High-Fidelity Models of Sense LincRNA-p21 *AluSx1* RNA. **Figure 6** presents the high-resolution, high-fidelity sense models that have good fitting with their SAXS envelope as demonstrated by low NSD values. **A** represents model 514; **B** represents model 1036; **C** represents model 1476; **D** represents model 1677; and **E** represents model 1794 which exhibit chemically probed secondary structures: left arm (Blue); 5′-junction (Green); three-way junction (Yellow), right arm (Magenta), and the 3′-adenyl tail (Cyan). Terminal nucleotides are displayed as: 5′nt (Red, Sphere Modelled) and 3′nt (Lime Green, Sphere Modelled). Models are rotated along their x-axis by 90° as indicated by the inset. A flexible, single-stranded linker sequence is represented centrally (Orange).

**Figure 7** similarly presents the high-fidelity, high-resolution structures of antisense LincRNAp21 *AluSx1* which were modelled using previous chemically probed secondary structure predictions using SimRNA v3.20 [21]. They exhibit the left and right arms, and the 5′-three-way junctions. Both the left (Blue) and right (Magenta) arms of antisense LincRNA-p21 *AluSx1* project laterally in line with the x-axis. The identified 5′-uridyl tail (Cyan) consistently wraps around the left arm. Both the 3′-three-way junctions, and the identified 5′-junction are compacted centrally, with regions that project perpendicularly from the RNA body’s x-axis. The right arm of antisense LincRNA-p21 *AluSx1* does not retain the characteristic stem-loop head that sense LincRNA-p21 *AluSx1* has. An animated representation of **Figure 7C** is attached in **Supplementary Information** as **SM 2**.

**Figure 7:**
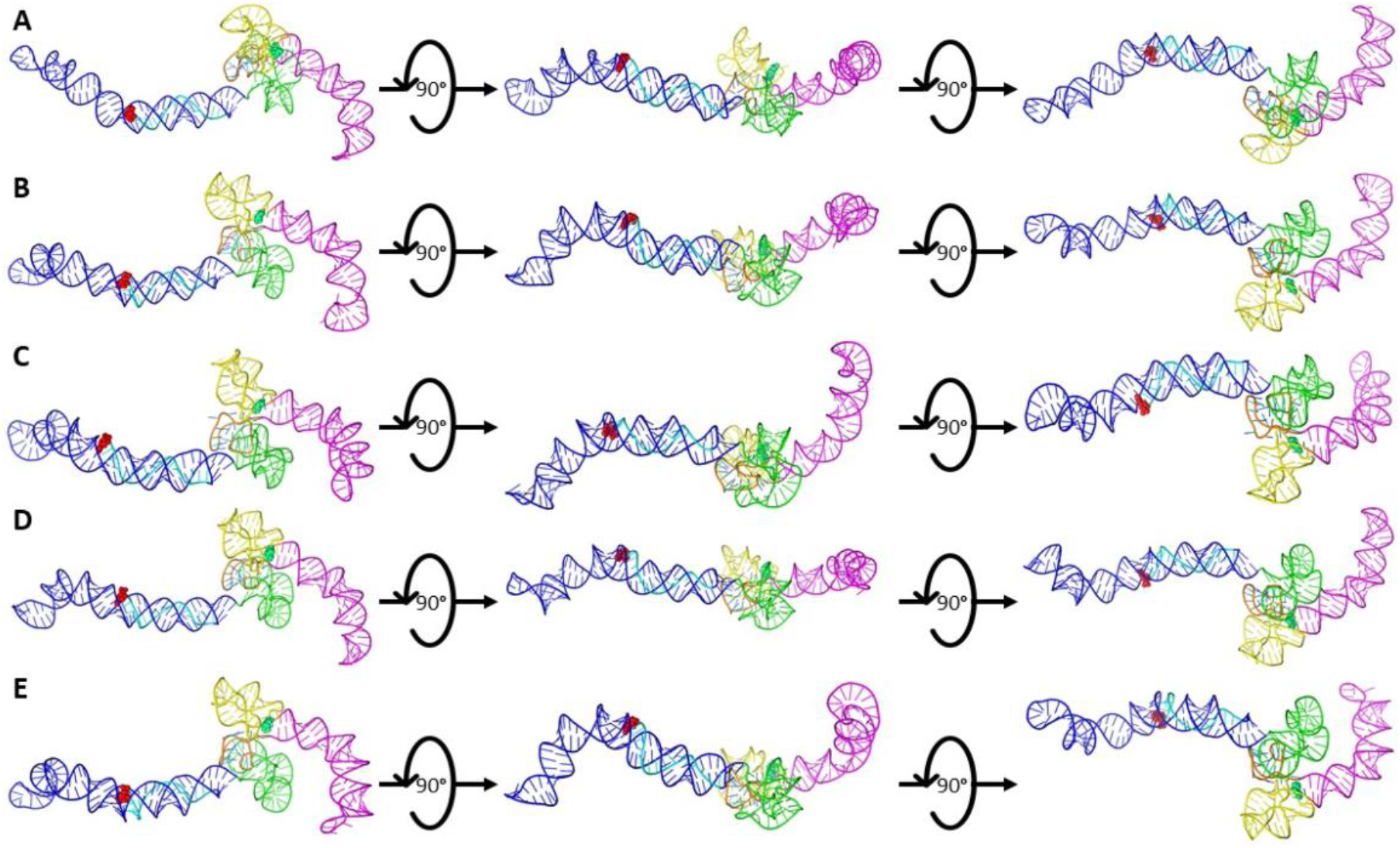
The SimRNA High-Resolution, High-Fidelity Models of Antisense LincRNA-p21 *AluSx1* RNA. **Figure 7** presents the high-resolution, high-fidelity sense models that have good fitting with their SAXS envelope as demonstrated by low NSD values. **A** represents model 66; **B** represents model 974; **C** represents model 1013; **D** represents model 1074; and **E** represents model 1417 which exhibit chemically probed secondary structures: left arm (Blue); 5′-junction (Green); three-way junction (Yellow), right arm (Magenta), and the 5′-uridyl tail (Cyan). Terminal nucleotides are displayed as: 5′nt (Red, Sphere Modelled) and 3′nt (Lime Green, Sphere Modelled). Models are rotated along their x-axis by 90° as indicated by the inset. A flexible, single-stranded linker sequence is represented centrally (Orange).

After performing DAMSUP, the high-fidelity, high-resolution models were visually inspected in terms of their alignment with the low-resolution SAXS envelope from DAMFILT and were represented in **Figures 8** and **9** for sense and antisense *AluSx1* RNA respectively. **Figure 8** displays a general agreement with the sense SAXS envelope overlaid by their high-fidelity, high resolution SimRNA models. Both the left and right arms when superimposed fit neatly within the protruding two bulges of the SAXS envelope. Minor disagreement occurs for the right arm’s stem-loop head which appears to be excluded from the SAXS structure, which is similar for the tip of the Left arm. Both the 5′-junction and 3′-three-way junction exhibits considerable overlap with the SAXS low-resolution structure when superimposed. **Figure 9** shows a general agreement with the antisense SimRNA models with their respective SAXS low-resolution structures. The left arm, 5′-junction, and 3′-three-way junctions are secondary structures that exhibit the highest relative overlap with the SAXS envelope and occur within the central bulge and right-most protrusion. The right arm (Magenta) depicts relatively lower agreement, with its tip and core regions exposed and externalised from the SAXS envelope. This is primarily confined to the left bulge, however, which indicates a lack of adequate fitting. Two animated representations of one of the sense (**Figure 8B**) and antisense (**Figure 9C**) models are attached in **Supplementary Information** as **SM 3** and **SM 4** respectively for demonstrating the lowest NSD values 1.073 and 1.156 respectively of the five selected models.

**Figure 8:**
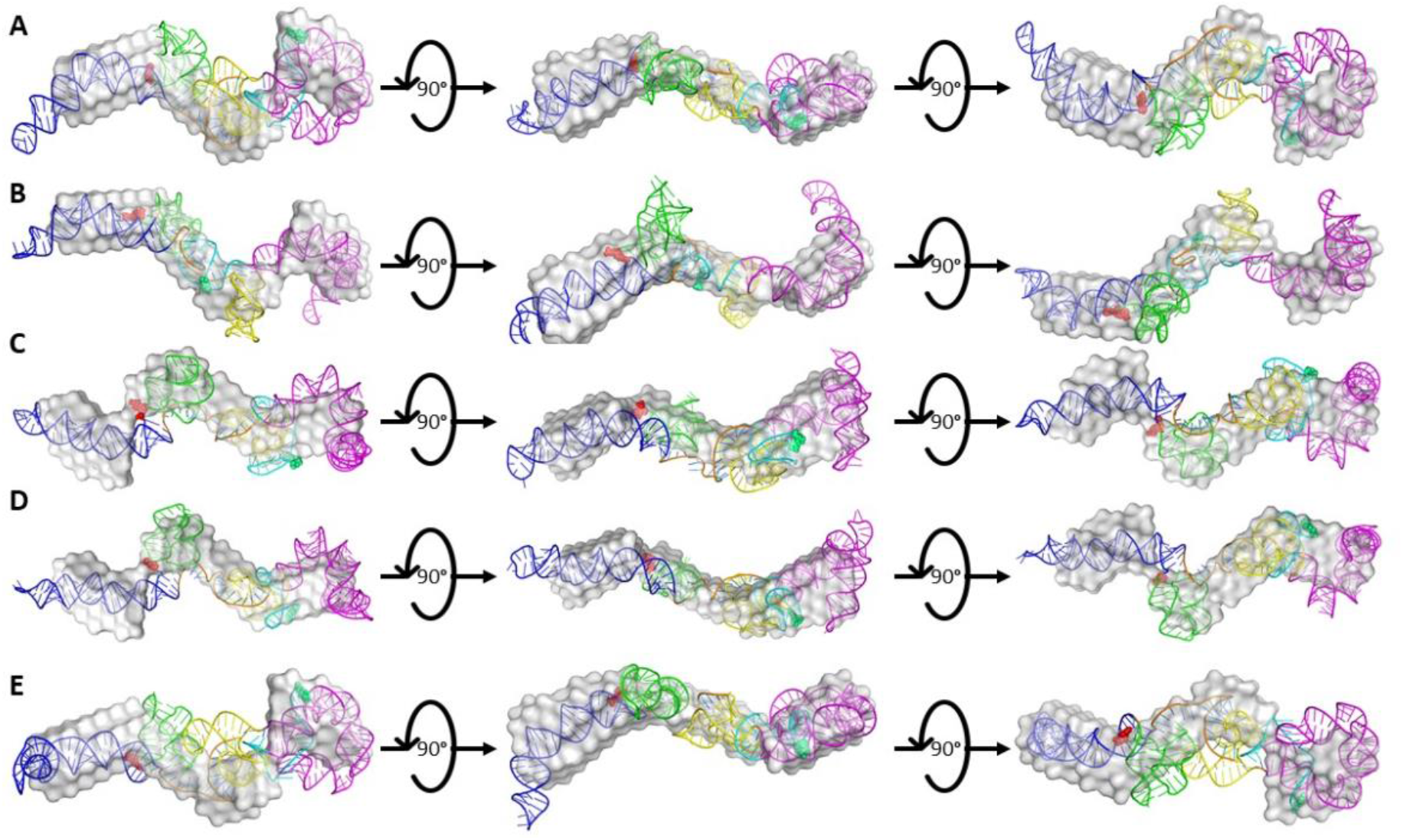
Superimposed Overlays of Sense SAXS Envelopes with their High-Resolution, High-Fidelity SimRNA Models. **Figure 8** represents the combined overlays of the sense LincRNA-p21 *AluSx1* RNA envelope with their top high-resolution, SimRNA models: (**A**) 514; (**B**) 1036; (**C**) 1476; (**D**) 1677; and (**E**) 1794. Overall, SAXS envelopes indicate a general agreement with computationally generated structures, showing high overlap of the extended molecule with the major secondary structures identified using chemical probing techniques.

**Figure 9:**
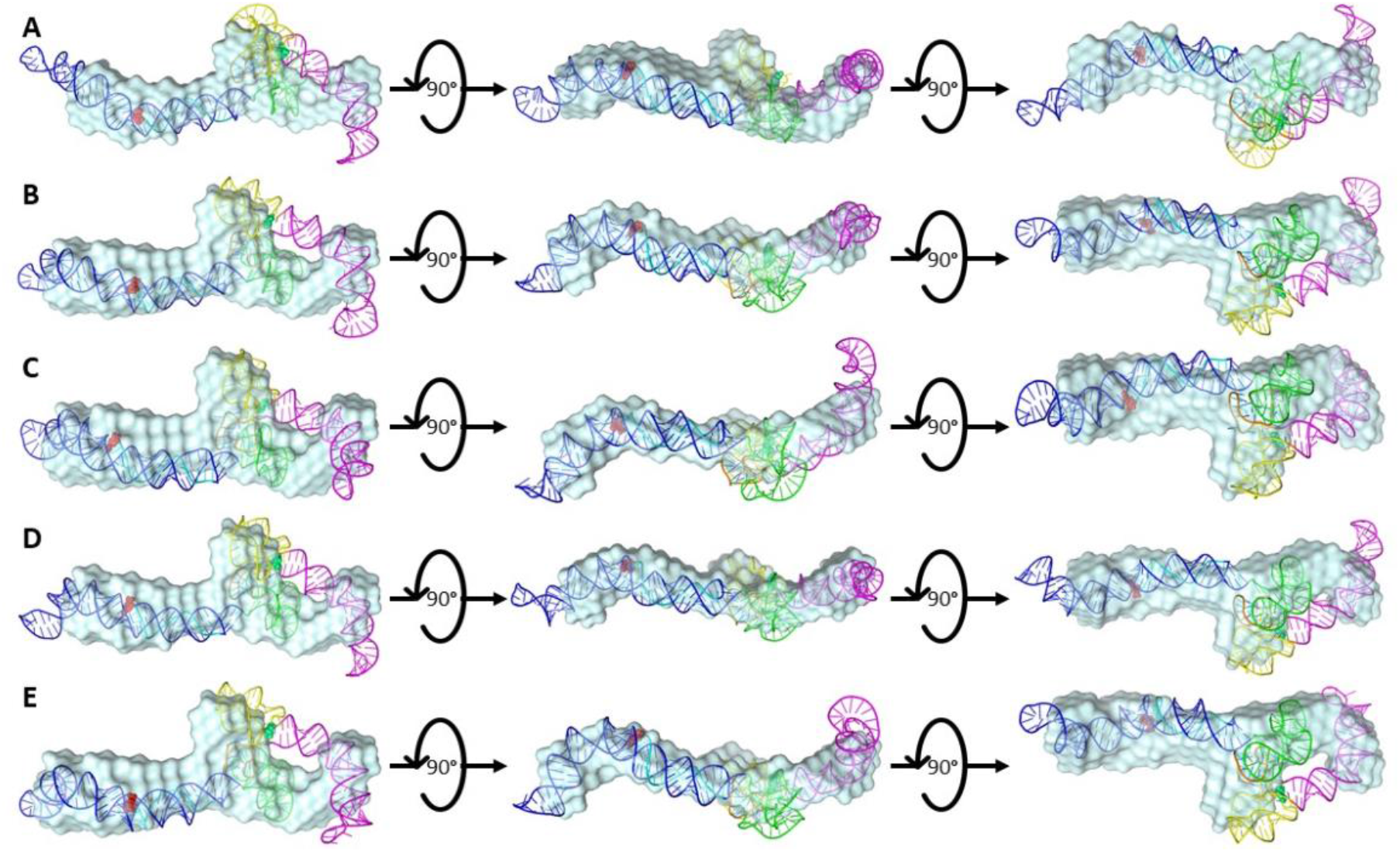
Superimposed Overlays of Antisense SAXS Envelopes with their High-Resolution, High-Fidelity SimRNA Models. **Figure 9** represents the combined overlays of the antisense LincRNA-p21 *AluSx1* RNA envelope with their top high-resolution, SimRNA models: (**A**) 66; (**B**) 974; (**C**) 1013; (**D**) 1074; and (**E**) 1417. Overall, SAXS envelopes indicate a general agreement with computationally generated structures, showing high overlap of the extended molecule with the major secondary structures identified using chemical probing techniques. However, there is a slight overhang present with the right arm (Magenta) which has an area excluded from overlapping with the SAXS envelope.

## 4.0 Discussion

RNA secondary structure is especially important in defining an RNA molecule’s roles and functions [72, 73]. These structures can be studied using a variety of techniques including secondary structure probing methods such as SHAPE, Nuclear Magnetic Resonance (NMR) spectroscopy, comparative sequence alignment and analysis, or three-dimensional structure predictions software for conceptualising higher order complexes [74–78]. Identifying functional structural elements of RNA is especially important when observing the noncoding elements of the genome, whose significant contribution of noncoding RNA (ncRNA) play important regulatory roles in complex organisms. LincRNA-p21 is one such important regulatory lncRNA that is directly targeted by p53 in response to DNA damage. Being a transcriptional repressor in the p53 pathway, understanding LincRNA-p21’s secondary structure is a first step evaluation towards how it binds to and modulates hnRNP-K localisation which is a process that ultimately triggers apoptosis in DNA damaged or cancerous cells.

The primary intent of our work is to obtain the high-resolution details of LincRNA-p21 *AluSx1* RNA by combining the previously determined secondary structure of the Inverted Repeat transposable elements determined by Chillón and Pyle with SAXS structure determination and computational modelling [21]. The previous study observed that the two isoforms of hLincRNA-p21 contain IR *Alu* elements, with the sense *AluSx1* located at positions 2589-2895 and the antisense *AluSx1* located at positions 1351-1651 of the TP53COR1 gene [21]. Since these RNAs are transcribed bidirectionally, observation of the secondary structure details of both the sense and antisense *AluSx1* was studied. **Figure 1** outlines our multifaceted process for biophysically characterising molecules using SEC-SAXS, SEC-MALS, SV-AUC, and SimRNA from chemically probed secondary structure determinations. SEC-MALS and AUC act as orthogonal quality control and validation techniques when combined with PAGE [79].

We initially started by transcribing sense and antisense *AluSx1* RNAs, which were purified to homogeneity as indicated by **Figure 2A**. Both RNA displays multi-modal peaks with a primary peak representative of the monomeric RNA species. Fractions along the primary peak were taken and analysed by 6M urea PAGE, which showed relatively pure and pronounced RNA bands around 300 bp indicating that most of the species is the primary RNA of interest (**Figure 2B**). Subsequent urea PAGEs were performed, and each resulted in an inconclusive answer to RNA homogeneity. We therefore turned to SV-AUC to determine if the RNA samples were heterogenous as indicated by urea-PAGE, or homogenous enough for further characterisation. As indicated in **Figure 2C, D**, both sense and antisense *AluSx1* RNA adopted a single, major species in solution when denatured by 6M urea. Any minor species had negligible concentrations and reflected noise contributions. The SV-AUC experiments showed no evidence for RNA degradation or aggregation. The MW values of sense and antisense *AluSx1* RNAs determined by AUC were less than 5% different (**Table 1**). AUC experiments were carried out in the presence of 6M urea as a denaturant, so we needed to assess each RNA in non-denaturing conditions. We therefore utilized SEC-MALS to determine absolute molecular weight in solution [41, 80]. Each RNA eluted as a singular, tight, and gaussian distribution elution profile evident in **Figure 3A**. Comparatively to AUC results, the RNA also exists monomerically in HFB as indicated by their uniform MW demonstrated in **Figures 3B/C**. The sense RNA molecular weight resulted in 99.24 ± 0.01, virtually identical to the theoretical molecular weight of 99.42 kDa. Antisense RNA also resulted in a very similar molecular weight of 94.52 ± 3.71 to the theoretical value of 89.54 kDa. These results suggest the RNA was acceptable to undergo SEC-SAXS and three-dimensional structure determination since SAXS requires highly homogenous samples.

SAXS is an ideal technique to determine low-resolution, three-dimensional structures of molecules in their native state and was used to investigate the solution conformations of both RNAs [59, 81, 82]. SAXS data is derived from the elastic scattering and detection of high-energy photons from the electrons of an irradiated sample which can be used to examine the sample’s electron density and overall structure [81]. The process of SAXS involves a solubilised sample exposed to a collimated monochromatic x-ray beam which scatters as it comes in contact with the sample’s molecules in question [82]. The resulting scattered x-rays can be measured based upon their intensity and altered angle verses incident and scattered beams. The solvent is measured independently to be subtracted as background noise and to remove interfering signals. Data analysis follows *ab initio* shape determination which takes the scattering pattern and represents them by finite volume elements (beads). Monte-Carlo type algorithms are employed to fit the experimental data at first using random configurations. DAMMIN and DAMMIF are *ab initio* programs that can be utilised to develop compact and interconnected bead models that illustrate the overall structure of the RNA in question [59, 83]. SAXS sampling is relatively easy and merely includes the purification of a sample (using SEC or filtering) and its subsequent exposure to an X-ray source. Such samples have the benefit of being observed in their near-physiological conditions without having to undergo artificial adjustments such as complex labelling, crystallisation, or cryogenic freezing. SAXS, however, has some limitations, namely its lower resolution compared to Cryo-Electron Microscopy or X-ray Crystallography causing ambiguity in discerning structural elements from scattering profiles [84]. Additionally, x-rays can damage samples but can be minimised with buffer protectants like glycerol [85].

Overall, scattering intensities were acceptable even in low-angle regions as shown in **Figure 4A**. The linear regression of the sense and antisense *AluSx1* samples in **Figure 4B** portrays the intensities within the defined low-q^2^ range as being linear with the absence of upward curves, illustrating monodispersity and the absence of attractive or repulsive interactions between scatterers [69, 86, 87]. Guiner R_g_ approximation resulted in 60.87 ± 0.85 and 59.07 ± 0.15 for sense and antisense *AluSx1* RNAs respectively. These Rg approximations are consistent with an elongated RNA molecule [88, 89]. Furthermore, the relative foldedness of sense and antisense RNAs can be deduced from the dimensionless Kratky plot in **Figure 4C**. The Kratky plot depicts both sense and antisense RNAs as being elongated but relatively folded, like other ncRNA [36, 90]. Additionally, the Guinier R_g_ and the real-space R_g_ are close with less than 1.5% difference. **Figure 4D** represents both RNA’s distance distribution functions and are non-Gaussian, further consistent with extended molecules. Globular molecules will generate a Gaussian-like P(r) distributions which is not demonstrated in **Figure 4D** further justifying its extended shape [70]. Using the P(r) distribution, molecules can also be described on their overall shape and symmetry to confirm solution folding acquired from the dimensionless Kratky plot [70]. Generally, globular molecules will display a bell-shaped curve with a maximum at approximately D_max_/2 while elongated molecules retain non-Gaussian, asymmetrical distributions with a maximum at smaller distances which appear as shoulders. For elongated molecules, this distribution will correspond to the radius of the cross section which will generally be illustrated by a tailing of the profile at larger distances [36, 91]. This is present for both sense and antisense *AluSx1* RNA whereby their respective D_max_/2 do not demonstrably produce an even bell-curved but rather lead into right-leaning tails reinforcing the overall elongated shape established by **Figure 4C**. Sense and antisense adopt maximum distance towards 185.0 Å and 180.7 Å respectively. Both RNAs result in similar D_max_ measurements which is expected of two RNA of similar lengths, with the sense RNA being larger than the antisense RNA. These size difference can be attributed both to the RNAs’ folding properties, and the 307 nt vs 280 nt lengths of the sense and antisense sequences as derived from Chillón and Pyle [21].

SAXS analysis of both the sense and antisense RNAs provided structures with noticeably consistent features, including a left and right bulge for the sense RNA and a left and central bulge, and right protrusion for the antisense RNA (**Figure 5**). Use of the standard HFB was important since human LincRNA-p21 folds at near physiological concentrations at around 5mM MgCl_2_ with maximum compaction at 15mM MgCl2 [21]. These features are observed to overlap with predicted features seen in the secondary structure analysis by Chillion and Pyle [21]. These features, while they can be seen in the SAXS structures, were arbitrarily based on structure orientation because directionality cannot be determined from these models. Therefore, we needed to not only computationally derive higher resolution models for clarity, but also directionality.

When analysing the high-resolution computational models derived from the secondary structure determinations, a straitened selection process was needed to screen for models that match the experimentally determined low-resolution structures. Therefore, HYDROPRO was employed to compute the hydrodynamic properties of sense and antisense *AluSx1* rigid macromolecules from their atomic-level structure [92–95]. HYDROPRO’s calculation comprise the basic hydrodynamic properties including the translational diffusion coefficient, sedimentation coefficient, intrinsic viscosity, and relaxation times, and can additionally provide the radius of gyration [61]. Incidentally, HYDROPRO can be used as another orthogonal selection process against computationally generated models that don’t fit the solutions scattering data. It was employed to minimise the ~3080 models which could subsequently be fit using DAMSUP based on the solution scattering range of D_max_ and R_g_. Consequently, observing HYDROPRO merely as a fitting tool narrowed the selection of potential models to five.

As detailed in **Figures 6 and 7**, sense and antisense *AluSx1* RNA folds into a double-stranded RNA molecule that appear to be consistent with Chillón and Pyle’s secondary structure determinations. No previous attempts at secondary structure determination of *AluSx1* has been performed before Chillón and Pyle, although alternative techniques such as *in vivo* click (ic) SHAPE has been suggested as a technique to study the secondary structure of LncRNAs natively outside of the *in vitro* techniques applied [96–98]. Human LincRNA-p21 is itself a linear, single exon lncRNA which contains IR Alu repeats such as *AluSx1* [21]. Both the sense and antisense *AluSx1* RNAs were determined to comprise a left and right Arm with both arms connected by a single-stranded region. The 5′ domain of each arm is characterised by a central three-way junction while the 3′ domain is characterised by a long stem-loop. These subsequently form independent structural domains which contribute to a variety of core functions such as human LincRNA-p21’s nuclear localisation following cell stress and DNA damage events [20]. They suggested that the 5′-end of LincRNA-p21 interacts with hnRNP-K which could also regulate its nuclear localisation in conjunction with its *AluSx1* inverted repeat elements.

Our low-resolution, three-dimensional structure of sense LincRNA-p21 *AluSx1* RNA (**Figure 8**) exhibits what appears to be two dsRNA arms that fold helically into the otherwise elongated, but radially compact model. When overlaid with the SAXS envelope, considerable overlap is present between the high-fidelity, high-resolution models. Both the 5′-junction and 3′-three-way junctions are tightly localised in the centre of the RNA body, while the left and right arms branch outwards. This likely forms a binding pocket or coordination site for hnRNP-K’s one of three KH RNA/DNA binding domains which is involved in eliciting transcription regulation in the nucleus [99–101]. Antisense LincRNA-p21 *AluSx1* RNA is again represented as a dsRNA molecule with both the 5′-junction and 3′-three-way junctions being compressed around the central bulge of the SAXS envelope (**Figure 9**). The RNA molecule itself is mostly extended with the left and right arms projecting outward which likely contributes towards interacting with hnRNP-K RNA binding domains. Specifically, sequences enriched with Alu repeats that contained three stretches of at least six pyrimidines (C/T) and matching the consensus sequence RCCTCCC (R=A/G) derived from a SINE-derived nuclear RNA LOcalisation (SIRLOIN) element is found to interact with hnRNP-K to increase lncRNA nuclear enrichment [100, 102]. Three are apparent within the antisense *AluSx1*, with two on each junction and one on the left arm suggesting an interaction with the hnRNP-K’s three KH RNA-binding domains. This has been shown to be reliant on triplets of C/T regions [103].

The flexibility and tendency towards being unfolded for single-stranded regions creates difficulty in visualising an RNA structure absolutely which would require additional SAXS optimisation methods to represent [104, 105]. However, the impact of ssRNA regions is limited to the 5′-adenyl/3′-uridyl tails and central linker, with the former generally wrapping around its most proximate arm. Despite this, **Figures 6** and **7** show multiple high-resolution models which were available within the constraints set by the solution scattering data. Observing **Supplementary Movie 5**, portraying the superimposed sense *AluSx1* models, we identified that the five models generally overlapped at the left arm, the 5′-junctions, and 3′-three-way junctions exhibiting minor conformational differences that manifested largely at flex points in the left arm. Major differences occurred in the right arm whose inward folding deviated between each individual models. This lowered confidence into an absolute, single-representative model for the sense *AluSx1* structure. **Supplementary Movie 6** similarly shows the superimposed antisense *AluSx1* models, however, they exhibited clear visible overlap and uniformity throughout the four major secondary structure elements. This indicates minimal conformational differences, with the models being relatively indistinguishable from each other suggesting that all five models are equally likely given their close R_g_ and D_max_ parameters to the solution scattering data. Agreement of solution structure and computational models for sense LincRNA-p21 *AluSx1* is evident by high overlaps and tight fitting. However, tangible exclusions are present for antisense LincRNA-p21 *AluSx1* RNA caused by a lack of overlap with the right arm and the tip of the left arm. This can potentially be attributed to the difference in length between the solution scattering sequence (280 nts) and the secondary structure prediction sequence (301 nts). The difference can be explained by construct synthesis whose sequence lacked 41 nts at the 3′-end but encompassed the remaining 80% of the secondary structure sequence predicted which still comprises the core and majority of the SAXS structure (**Supplementary Information)**. Nevertheless, both structures detail the presence of the previously identified secondary structures and their orientations. Despite this, the best fitting models that uniquely overlapped with their solution scattering three-dimensional structures are **Figures 8B** and **9C** for sense and antisense *AluSx1* due to their low NSD values. Altogether, the models are not an ideal, true fits which would constitute a zero-NSD value and is largely the result of systematic differences [63, 106]. The discernible difference in antisense *AluSx1* is accounted by the deleted 3′-end while overall differences can be accounted to SAXS bead to atom approximations within SUPCOMB itself.

Nevertheless, applying both solution scattering techniques and coarse-grained computational modelling reveals that LincRNA-p21 Alu Inverted Repeats do not adopt completely stable, single-representative structures due in part to the conformational flexibility present in their respective DAMMIN and SimRNA models. Both the sense and antisense *AluSx1* RNAs adopt multiple conformations, however, they closely approximate into a generally similar, single-representative structure with mild conformational and structural differences when averaged. The main shape – one that is elongated, asymmetrical, with regions that encompass the main left and right arms, and three-way junction – is uniform and preserved throughout solution scattering and computational fitting techniques. Consequently, applying a combination of SAXS and computationally generated tertiary structure models concertedly determined appropriate representations of the LincRNA-p21 *AluSx1* Inverted Repeats.

## 5.0 Conclusion

We have demonstrated that chemical probing techniques involved in RNA secondary structure predictions such as SHAPE can be combined using a multifaceted biophysical approach involving SAXS, AUC, SEC-MALS and computational RNA modelling. Although generally averaged, the three-dimensional structures of sense and antisense LincRNA-p21 *AluSx1* IRs provided from **SM3** and **SM4** respectively represent a current understanding of their overall structure. Naturally, higher resolution can be achieved using NMR and single particle Cryo-Electron Microscopy methods, employing similarly computational techniques to acquire as low as 3.7 Å resolution on very small (~100 nt) RNA [78, 107, 108]. This would be to expand on structure correlation between crystal or solution scattering methods and computational ones to overcome the heterogeneous and flexible nature of RNAs. Nevertheless, three-dimensional models are important in confirming secondary structure motifs that are predicted through probing techniques. By expanding the structure to include three-dimensional native state folding, essential regulatory RNAs involved in apoptosis and tumour suppression in cancer cells can be effectively visualised to identify functional domains and potential RNA-Protein binding regions.

## Acknowledgements and Funding Support

M.H.D and M.G. are supported by the NSERC Discovery grant awarded to T.R.P. M.H.D is supported by the South Alberta Light Horse Regimental Foundation, The King’s Own Calgary Regiment (RCAC) Funds Foundation, and the Canadian Armed Forces Individual Learning Plan and Self-Development Program. M.D.B. is supported by Alberta Innovates grant to T.R.P. T.M. is supported by a Natural Sciences and Engineering Research Council (NSERC) PGS-D award. A.H. is supported by the NSERC CGS-M award. T.R.P. is a Canada Research Chair in RNA and Protein Biophysics. Infrastructure support to T.R.P. and B.D. was provided by the Canada Foundation for Innovation and NSERC RTI Grants. This work was supported by the Canada 150 Research Chairs program (B.D., C150-2017-00015), the Canada Foundation for Innovation (B.D., CFI-37589), the National Institutes of Health (B.D., 1R01GM120600) and the Canadian Natural Science and Engineering Research Council (B.D., DG-RGPIN-2019-05637). UltraScan supercomputer calculations were supported through NSF/XSEDE grant TG-MCB070039N (to B.D).The SAXS data collection was supported by DIAMOND Synchrotron, UK. We thank all the B21 beamline scientists at DIAMOND for their continual support.

## Author Contribution

M.H.D., M.G., and A.F. prepared RNAs and purified them. M.H.D., M.D.B., and T.R.P. processed SAXS data. T.M. and M.H.D. collected and processed SEC-MALS data. A.H. collected AUC data. A.H., B.D., and M.H.D. processed AUC data. M.W. performed computational RNA modelling. M.T.W., M.H.D., and T.R.P. analysed computational models. M.H.D. developed initial drafts of manuscript. All authors reviewed and edited manuscript.

## References

1. Rufini, A., et al., Senescence and aging: the critical roles of p53. Oncogene, 2013. 32(43): p. 5129–43.

2. Amaral, J.D., et al., The role of p53 in apoptosis. Discov Med, 2010. 9(45): p. 145–52.

3. Treviño, V., E. Martínez-Ledesma, and J. Tamez-Peña, Identification of outcome-related driver mutations in cancer using conditional co-occurrence distributions. Sci Rep, 2017. 7: p. 43350.

4. Catana, C.-S., D. Gulei, and I. Berindan - Neagoe, New insights into the role of non-coding RNAs as transcriptional targets of p53. Endogenous locus-driven H-Ras G12V expression induces senescence-like phenotype in primary fibroblasts of the Costello syndrome mouse model, 2017: p. 43–49.

5. Hafner, A., et al., The multiple mechanisms that regulate p53 activity and cell fate. Nat Rev Mol Cell Biol, 2019. 20(4): p. 199–210.

6. Harris, S.L. and A.J. Levine, The p53 pathway: positive and negative feedback loops. Oncogene, 2005. 24(17): p. 2899–908.

7. Appella, E. and C.W. Anderson, Post-translational modifications and activation of p53 by genotoxic stresses. Eur J Biochem, 2001. 268(10): p. 2764–72.

8. Levine, A.J., p53: 800 million years of evolution and 40 years of discovery. Nat Rev Cancer, 2020. 20(8): p. 471–480.

9. Oliner, J.D., et al., Oncoprotein MDM2 conceals the activation domain of tumour suppressor p53. Nature, 1993. 362(6423): p. 857–60.

10. Haupt, Y., et al., Mdm2 promotes the rapid degradation of p53. Nature, 1997. 387(6630): p. 296–9.

11. Yin, Y., et al., p53 Stability and activity is regulated by Mdm2-mediated induction of alternative p53 translation products. Nat Cell Biol, 2002. 4(6): p. 462–7.

12. Gu, W. and R.G. Roeder, Activation of p53 sequence-specific DNA binding by acetylation of the p53 C-terminal domain. Cell, 1997. 90(4): p. 595–606.

13. Guttman, M., et al., Chromatin signature reveals over a thousand highly conserved large non-coding RNAs in mammals. Nature, 2009. 458(7235): p. 223–7.

14. Jin, S., et al., p53-targeted lincRNA-p21 acts as a tumor suppressor by inhibiting JAK2/STAT3 signaling pathways in head and neck squamous cell carcinoma. Mol Cancer, 2019. 18(1): p. 38.

15. Wang, X., et al., Long intragenic non-coding RNA lincRNA-p21 suppresses development of human prostate cancer. Cell Prolif, 2017. 50(2).

16. Tran, U.M., et al., LincRNA-p21 acts as a mediator of ING1b-induced apoptosis. Cell Death Dis, 2015. 6(3): p. e1668.

17. Chen, S., et al., LincRNa-p21: function and mechanism in cancer. Med Oncol, 2017. 34(5): p. 98.

18. Tang, S.S., B.Y. Zheng, and X.D. Xiong, LincRNA-p21: Implications in Human Diseases. Int J Mol Sci, 2015. 16(8): p. 18732–40.

19. Kesheh, M.M., S. Mahmoudvand, and S. Shokri, Long noncoding RNAs in respiratory viruses: A review. Rev Med Virol, 2021: p. e2275.

20. Huarte, M., et al., A large intergenic noncoding RNA induced by p53 mediates global gene repression in the p53 response. Cell, 2010. 142(3): p. 409–19.

21. Chillón, I. and A.M. Pyle, Inverted repeat Alu elements in the human lincRNA-p21 adopt a conserved secondary structure that regulates RNA function. Nucleic Acids Res, 2016. 44(19): p. 9462–9471.

22. Moumen, A., et al., hnRNP K: an HDM2 target and transcriptional coactivator of p53 in response to DNA damage. Cell, 2005. 123(6): p. 1065–78.

23. Sun, X., M.S.S. Haider Ali, and M. Moran, The role of interactions of long non-coding RNAs and heterogeneous nuclear ribonucleoproteins in regulating cellular functions. The Biochemical journal, 2017. 474(17): p. 2925–2935.

24. Baldassarre, A. and A. Masotti, Long non-coding RNAs and p53 regulation. Int J Mol Sci, 2012. 13(12): p. 16708–17.

25. Arcot, S.S., et al., Alu repeats: a source for the genesis of primate microsatellites. Genomics, 1995. 29(1): p. 136–44.

26. Batzer, M.A. and P.L. Deininger, Alu repeats and human genomic diversity. Nat Rev Genet, 2002. 3(5): p. 370–9.

27. Tajaddod, M., et al., Transcriptome-wide effects of inverted SINEs on gene expression and their impact on RNA polymerase II activity. Genome Biol, 2016. 17(1): p. 220.

28. Deininger, P.L. and M.A. Batzer, Alu repeats and human disease. Mol Genet Metab, 1999. 67(3): p. 183–93.

29. Payer, L.M., et al., Structural variants caused by Alu insertions are associated with risks for many human diseases. Proc Natl Acad Sci U S A, 2017. 114(20): p. E3984–e3992.

30. Polak, P. and E. Domany, Alu elements contain many binding sites for transcription factors and may play a role in regulation of developmental processes. BMC Genomics, 2006. 7: p. 133.

31. Novikova, I.V., S.P. Hennelly, and K.Y. Sanbonmatsu, Sizing up long non-coding RNAs: do lncRNAs have secondary and tertiary structure? Bioarchitecture, 2012. 2(6): p. 189–199.

32. Chillón, I. and M. Marcia, The molecular structure of long non-coding RNAs: emerging patterns and functional implications. Crit Rev Biochem Mol Biol, 2020. 55(6): p. 662–690.

33. Lusvarghi, S., et al., RNA secondary structure prediction using high-throughput SHAPE. J Vis Exp, 2013(75): p. e50243.

34. Boniecki, M.J., et al., SimRNA: a coarse-grained method for RNA folding simulations and 3D structure prediction. Nucleic Acids Res, 2016. 44(7): p. e63.

35. Conrad, T., et al., Maximizing transcription of nucleic acids with efficient T7 promoters. Commun Biol, 2020. 3(1): p. 439.

36. Mrozowich, T., et al., Nanoscale Structure Determination of Murray Valley Encephalitis and Powassan Virus Non-Coding RNAs. Viruses, 2020. 12(2).

37. Chillón, I., et al., Native Purification and Analysis of Long RNAs. Methods Enzymol, 2015. 558: p. 3-37.

38. Beckert, B. and B. Masquida, Synthesis of RNA by in vitro transcription. Methods Mol Biol, 2011. 703: p. 29–41.

39. Chen, Z. and Y. Zhang, Dimethyl sulfoxide targets phage RNA polymerases to promote transcription. Biochem Biophys Res Commun, 2005. 333(3): p. 664–70.

40. McKenna, S.A., et al., Purification and characterization of transcribed RNAs using gel filtration chromatography. Nat Protoc, 2007. 2(12): p. 3270–7.

41. Some, D., et al., Characterization of Proteins by Size-Exclusion Chromatography Coupled to Multi-Angle Light Scattering (SEC-MALS). J Vis Exp, 2019(148).

42. Pam Wang, R.A., Michelle Chen, and Kristine Legaspi, AN1616: SEC-MALS Method for Characterizing mRNA Biophysical Attributes. Wyatt Technologies Moderna Therapeutics 2020.

43. Pam Wang, R.A., Michelle Chen, and Kristine Legaspi, SEC-MALS Method for Characterizing mRNA Biophysical Attributes. LCGC 2020.

44. Patel, T.R., et al., Structural studies of RNA-protein complexes: A hybrid approach involving hydrodynamics, scattering, and computational methods. Methods, 2017. 118-119: p. 146–162.

45. Wyatt, P.J., Measurement of special nanoparticle structures by light scattering. Anal Chem, 2014. 86(15): p. 7171–83.

46. Wyatt, P.J., Measuring nanoparticles in the size range to 2000 nm. Journal of nanoparticle research : an interdisciplinary forum for nanoscale science and technology, 2018. 20(12): p. 322–322.

47. Demeler, B. and G.E. Gorbet, Analytical Ultracentrifugation Data Analysis with UltraScan-III, in Analytical Ultracentrifugation: Instrumentation, Software, and Applications, S. Uchiyama, et al., Editors. 2016, Springer Japan: Tokyo. p. 119–143.

48. Demeler, B., Methods for the design and analysis of sedimentation velocity and sedimentation equilibrium experiments with proteins. Curr Protoc Protein Sci, 2010. Chapter 7: p. Unit 7.13.

49. Brookes, E., W. Cao, and B. Demeler, A two-dimensional spectrum analysis for sedimentation velocity experiments of mixtures with heterogeneity in molecular weight and shape. Eur Biophys J, 2010. 39(3): p. 405–14.

50. Brookes, E.H. and B. Demeler, Parsimonious regularization using genetic algorithms applied to the analysis of analytical ultracentrifugation experiments, in Proceedings of the 9th annual conference on Genetic and evolutionary computation. 2007, Association for Computing Machinery: London, England. p. 361–368.

51. Demeler, B. and E. Brookes, Monte Carlo analysis of sedimentation experiments. Colloid and Polymer Science, 2008. 286(2): p. 129–137.

52. Cowieson, N.P., et al., Beamline B21: high-throughput small-angle X-ray scattering at Diamond Light Source. J Synchrotron Radiat, 2020. 27(Pt 5): p. 1438–1446.

53. Manalastas-Cantos, K., et al., ATSAS 3.0: expanded functionality and new tools for small-angle scattering data analysis. J Appl Crystallogr, 2021. 54(Pt 1): p. 343–355.

54. Panjkovich, A. and D.I. Svergun, CHROMIXS: automatic and interactive analysis of chromatography-coupled small-angle X-ray scattering data. Bioinformatics, 2018. 34(11): p. 1944–1946.

55. Putnam, C.D., Guinier peak analysis for visual and automated inspection of small-angle X-ray scattering data. J Appl Crystallogr, 2016. 49(Pt 5): p. 1412–1419.

56. Burke, J.E. and S.E. Butcher, Nucleic acid structure characterization by small angle X-ray scattering (SAXS). Current protocols in nucleic acid chemistry, 2012. Chapter 7: p. Unit7.18–Unit7.18.

57. Semenyuk, A.V. and D.I. Svergun, GNOM– a program package for small-angle scattering data processing. Journal of Applied Crystallography, 1991. 24(5): p. 537–540.

58. Svergun, D.I., Determination of the regularization parameter in indirect-transform methods using perceptual criteria. Journal of Applied Crystallography, 1992. 25(4): p. 495–503.

59. Svergun, D.I., Restoring low resolution structure of biological macromolecules from solution scattering using simulated annealing. Biophys J, 1999. 76(6): p. 2879–86.

60. Volkov, V.V. and D.I. Svergun, Uniqueness of ab initio shape determination in small-angle scattering. Journal of Applied Crystallography, 2003. 36(3-1): p. 860–864.

61. Ortega, A., D. Amorós, and J. García de la Torre, Prediction of hydrodynamic and other solution properties of rigid proteins from atomic- and residue-level models. Biophys J, 2011. 101(4): p. 892–8.

62. Stasiewicz, J., et al., QRNAS: software tool for refinement of nucleic acid structures. BMC Structural Biology, 2019. 19(1): p. 5.

63. Kozin, M.B. and D.I. Svergun, Automated matching of high- and low-resolution structural models. Journal of Applied Crystallography, 2001. 34(1): p. 33–41.

64. Brosey, C.A. and J.A. Tainer, Evolving SAXS versatility: solution X-ray scattering for macromolecular architecture, functional landscapes, and integrative structural biology. Curr Opin Struct Biol, 2019. 58: p. 197–213.

65. Pérez, J. and P. Vachette, A Successful Combination: Coupling SE-HPLC with SAXS. Adv Exp Med Biol, 2017. 1009: p. 183–199.

66. O’Brien, D.P., et al., SEC-SAXS and HDX-MS: A powerful combination. The case of the calcium-binding domain of a bacterial toxin. Biotechnol Appl Biochem, 2018. 65(1): p. 62–68.

67. Gräwert, M., et al., Adding Size Exclusion Chromatography (SEC) and Light Scattering (LS) Devices to Obtain High-Quality Small Angle X-Ray Scattering (SAXS) Data. Crystals, 2020. 10: p. 975.

68. Rambo, R.P. and J.A. Tainer, Characterizing flexible and intrinsically unstructured biological macromolecules by SAS using the Porod-Debye law. Biopolymers, 2011. 95(8): p. 559–571.

69. Putnam, C.D., et al., X-ray solution scattering (SAXS) combined with crystallography and computation: defining accurate macromolecular structures, conformations and assemblies in solution. Q Rev Biophys, 2007. 40(3): p. 191–285.

70. Kikhney, A.G. and D.I. Svergun, A practical guide to small angle X-ray scattering (SAXS) of flexible and intrinsically disordered proteins. FEBS Lett, 2015. 589(19 Pt A): p. 2570–7.

71. Vadlamani, G., et al., The β-lactamase gene regulator AmpR is a tetramer that recognizes and binds the D-Ala-D-Ala motif of its repressor UDP-N-acetylmuramic acid (MurNAc)-pentapeptide. J Biol Chem, 2015. 290(5): p. 2630–43.

72. Vandivier, L.E., et al., The Conservation and Function of RNA Secondary Structure in Plants. Annual review of plant biology, 2016. 67: p. 463–488.

73. Chełkowska-Pauszek, A., et al., The Role of RNA Secondary Structure in Regulation of Gene Expression in Bacteria. International journal of molecular sciences, 2021. 22(15): p. 7845.

74. Lorenz, R., et al., Predicting RNA secondary structures from sequence and probing data. Methods, 2016. 103: p. 86–98.

75. Chen, J.L., S. Bellaousov, and D.H. Turner, RNA Secondary Structure Determination by NMR. Methods Mol Biol, 2016. 1490: p. 177–86.

76. Kenyon, J., L. Prestwood, and A. Lever, Current perspectives on RNA secondary structure probing. Biochem Soc Trans, 2014. 42(4): p. 1251–5.

77. Tahi, F., T.T.V. Du, and A. Boucheham, In Silico Prediction of RNA Secondary Structure. Methods Mol Biol, 2017. 1543: p. 145–168.

78. Barnwal, R.P., F. Yang, and G. Varani, Applications of NMR to structure determination of RNAs large and small. Archives of biochemistry and biophysics, 2017. 628: p. 42–56.

79. Patel, B.A., et al., Multi-angle light scattering as a process analytical technology measuring real-time molecular weight for downstream process control. mAbs, 2018. 10(7): p. 945–950.

80. Stetefeld, J., S.A. McKenna, and T.R. Patel, Dynamic light scattering: a practical guide and applications in biomedical sciences. Biophys Rev, 2016. 8(4): p. 409–427.

81. Gräwert, T.W. and D.I. Svergun, Structural Modeling Using Solution Small-Angle X-ray Scattering (SAXS). J Mol Biol, 2020. 432(9): p. 3078–3092.

82. Svergun, D.I., Small-angle X-ray and neutron scattering as a tool for structural systems biology. Biol Chem, 2010. 391(7): p. 737–43.

83. Franke, D. and D.I. Svergun, DAMMIF, a program for rapid ab-initio shape determination in small-angle scattering. J Appl Crystallogr, 2009. 42(Pt 2): p. 342–346.

84. Jeffries, C.M., et al., Small-angle X-ray and neutron scattering. Nature Reviews Methods Primers, 2021. 1(1): p. 70.

85. Rambo, R.P. and J.A. Tainer, Improving small-angle X-ray scattering data for structural analyses of the RNA world. Rna, 2010. 16(3): p. 638–46.

86. Grant, T.D., et al., The accurate assessment of small-angle X-ray scattering data. Acta Crystallogr D Biol Crystallogr, 2015. 71(Pt 1): p. 45–56.

87. Cantara, W.A., E.D. Olson, and K. Musier-Forsyth, Analysis of RNA structure using small-angle X-ray scattering. Methods, 2017. 113: p. 46–55.

88. Zhang, Y., et al., Long non-coding subgenomic flavivirus RNAs have extended 3D structures and are flexible in solution. EMBO Rep, 2019. 20(11): p. e47016.

89. Cantero-Camacho, Á., et al., Three-dimensional structure of the 3’X-tail of hepatitis C virus RNA in monomeric and dimeric states. Rna, 2017. 23(9): p. 1465–1476.

90. Nelson, C.R., et al., Human DDX17 Unwinds Rift Valley Fever Virus Non-Coding RNAs. Int J Mol Sci, 2020. 22(1).

91. Choi, K.H. and M. Morais, Use of small-angle X-ray scattering to investigate the structure and function of dengue virus NS3 and NS5. Methods in molecular biology (Clifton, N.J.), 2014. 1138: p. 241–252.

92. Carrasco, B. and J. García de la Torre, Hydrodynamic properties of rigid particles: comparison of different modeling and computational procedures. Biophys J, 1999. 76(6): p. 3044–57.

93. Fernandes, M.X., et al., Calculation of hydrodynamic properties of small nucleic acids from their atomic structure. Nucleic Acids Res, 2002. 30(8): p. 1782–8.

94. García De La Torre, J., M.L. Huertas, and B. Carrasco, Calculation of hydrodynamic properties of globular proteins from their atomic-level structure. Biophys J, 2000. 78(2): p. 719–30.

95. Garcia de la Torre, J.G. and V.A. Bloomfield, Hydrodynamic properties of complex, rigid, biological macromolecules: theory and applications. Q Rev Biophys, 1981. 14(1): p. 81–139.

96. Feyder, M. and L.A. Goff, Investigating long noncoding RNAs using animal models. J Clin Invest, 2016. 126(8): p. 2783–91.

97. Sahu, A., U. Singhal, and A.M. Chinnaiyan, Long noncoding RNAs in cancer: from function to translation. Trends in cancer, 2015. 1(2): p. 93–109.

98. Spitale, R.C., et al., Structural imprints in vivo decode RNA regulatory mechanisms. Nature, 2015. 519(7544): p. 486–90.

99. Bomsztyk, K., O. Denisenko, and J. Ostrowski, hnRNP K: one protein multiple processes. Bioessays, 2004. 26(6): p. 629–38.

100. Xu, Y., et al., New Insights into the Interplay between Non-Coding RNAs and RNA-Binding Protein HnRNPK in Regulating Cellular Functions. Cells, 2019. 8(1): p. 62.

101. Makeyev, A.V. and S.A. Liebhaber, The poly(C)-binding proteins: a multiplicity of functions and a search for mechanisms. Rna, 2002. 8(3): p. 265–78.

102. Lubelsky, Y. and I. Ulitsky, Sequences enriched in Alu repeats drive nuclear localization of long RNAs in human cells. Nature, 2018. 555(7694): p. 107–111.

103. Paziewska, A., et al., Cooperative binding of the hnRNP K three KH domains to mRNA targets. FEBS Lett, 2004. 577(1-2): p. 134–40.

104. Meisburger, S.P., et al., Polyelectrolyte properties of single stranded DNA measured using SAXS and single-molecule FRET: Beyond the wormlike chain model. Biopolymers, 2013. 99(12): p. 1032–45.

105. Plumridge, A., S.P. Meisburger, and L. Pollack, Visualizing single-stranded nucleic acids in solution. Nucleic Acids Res, 2017. 45(9): p. e66.

106. Konarev, P.V., M.V. Petoukhov, and D.I. Svergun, Rapid automated superposition of shapes and macromolecular models using spherical harmonics. Journal of applied crystallography, 2016. 49(Pt 3): p. 953–960.

107. Kappel, K., et al., Accelerated cryo-EM-guided determination of three-dimensional RNA-only structures. Nat Methods, 2020. 17(7): p. 699–707.

108. Zhang, K., et al., Cryo-EM structure of a 40 kDa SAM-IV riboswitch RNA at 3.7 Å resolution. Nat Commun, 2019. 10(1): p. 5511.

